# Why can we detect lianas from space?

**DOI:** 10.1101/2021.09.30.462145

**Authors:** Marco D. Visser, Matteo Detto, Félicien Meunier, Jin Wu, Jane R. Foster, David C. Marvin, Peter M. van Bodegom, Boris Bongalov, Matheus Henrique Nunes, David Coomes, Hans Verbeeck, J. Antonio Guzmán Q, Arturo Sanchez-Azofeifa, Chris J. Chandler, Geertje M.F van der Heijden, Doreen S. Boyd, Giles M. Foody, Mark E.J. Cutler, Eben N. Broadbent, Shawn P. Serbin, Stefan Schnitzer, M. Elizabeth Rodríguez-Ronderos, Frank Sterck, José A. Medina-Vega, Steve Pacala

## Abstract

Lianas, woody vines acting as structural parasites of trees, have profound effects on the composition and structure of tropical forests, impacting tree growth, mortality, and forest succession. Remote sensing offers a powerful tool for quantifying the scale of liana infestation, provided the availability of robust detection methods. We analyze the consistency and global specificity of spectral signals from liana-infested tree crowns and forest stands, examining the underlying mechanisms. We compiled a database, including leaf reflectance spectra from 5424 leaves, fine-scale airborne reflectance data from 999 liana-infested canopies, and coarse-scale satellite reflectance data covering hectares of liana-infested forest stands. To unravel the mechanisms of the liana spectral signal, we applied mechanistic radiative transfer models across scales, corroborated by field data on liana leaf chemistry and canopy structure. We find a consistent liana spectral signature at canopy and stand scales across sites. This signature mainly arises at the canopy level due to direct effects of leaf angles, resulting in a larger apparent leaf area, and indirect effects from increased light scattering in the NIR and SWIR regions, linked to lianas’ less costly leaf construction compared to trees. The existence of a consistent global spectral signal for lianas suggests that large-scale quantification of liana infestation is feasible. However, because the traits identified are not exclusive to lianas, accurate large-scale detection requires rigorously validated remote sensing methods. Our models highlight challenges in automated detection, such as potential misidentification due to leaf phenology, tree life-history, topography, and climate, especially where the scale of liana infestation is less than a single remote sensing pixel. The observed cross-site patterns also prompt ecological questions about lianas’ adaptive similarities across environments, indicating possible convergent evolution due to shared constraints on leaf biochemical and structural traits.

**Open data statement:** Of the 17 datasets used, 10 are published and publicly accessible, with links provided in this submission (Appendix S1: Section S1). Upon acceptance, remaining seven datasets will be provided via Smithsonian’s Dspace. The open-source model code is available as R-package ccrtm (https://cran.r-project.org/web/packages/ccrtm/index.html) and on github (https://github.com/MarcoDVisser/ccrtm). Code will be archived in Zenodo should the manuscript be accepted for publication

## Introduction

Lianas are woody vines that act as structural parasites of trees and have a strong influence on forest dynamics (Visser et al. 2018b), decreasing tree growth and survival (Ingwell et al. 2010) and arresting forest succession (Tymen et al. 2016). Lianas are increasing in abundance in many forests (Schnitzer and Bongers 2011, Wright et al. 2015), raising concerns about the potential negative impact on carbon sequestration (Durán et al. 2013, van der Heijden et al. 2015). However, the spatial extent of this increase is uncertain, and the causes remain mostly unknown (Schnitzer 2015, Schnitzer et al. 2020). Remote sensing could provide a powerful tool to quantify this trend on a broad range of spatiotemporal scales and elucidate the climatic and bio-geographical factors contributing to liana proliferation (van der Heijden et al. 2022). Yet, as of today we do not know whether a globally distinct liana spectral-signal exists that could make such research possible.

Ecologists have utilized various remote sensing platforms and methods to detect lianas, including multispectral and hyperspectral sensors on drones, aircraft, and satellites (Foster et al. 2008, Tymen et al. 2016, Marvin et al. 2016, Li et al. 2018, Waite et al. 2019, Chandler et al. 2021, Kaçamak et al. 2022). These studies suggest that lianas have distinct detectable spectral signals at the canopy and stand scales. However, differentiating lianas and trees based on the spectral properties of leaves might depend on forest type, with detectable differences observed in dry forests but not in wet forests (Sánchez-Azofeifa et al. 2009a, Asner and Martin 2012, Guzmán Q. et al. 2018) suggesting that climatic factors influence the chemical and structural properties of liana leaves compared to their hosts (Asner and Martin 2012, Werden et al. 2018, Medina-Vega et al. 2021b).

The influence of climate indicates exceptions and contextual dependencies rather than a distinct global liana spectral signal. This raises doubts about the applicability of automated classifiers for quantifying liana abundances on large spatiotemporal domains. Data-driven automated classifiers can capture complex relationships but may struggle to generalize to scenarios outside the training data, particularly when crucial relationships are missing (Féret et al. 2019, Meyer and Pebesma 2022, Willard et al. 2022). The major challenge in robustly applying automated classification techniques is collecting training data representative of all conditions, which may not be feasible across large spatial domains with different climates or floristic compositions. Collecting representative training data in tropical forests is especially daunting due to confounders and high biodiversity. For instance, many leaf and canopy traits are known to vary systematically among plant groups with life history and leaf phenology, but differences can be small compared to the large variability across interspecific, intraspecific, phenotypic, and ontogenetic levels of natural vegetation (Castro-Esau et al. 2004, Kitajima et al. 2005, Zhang et al. 2006, Sánchez-Azofeifa et al. 2009a, Wu et al. 2018, Werden et al. 2018, Detto and Xu 2020).

At a more fundamental level, pure data-driven automated classifiers do not explain the underlying reasons responsible for a plant spectral signal. Alternatively, a process-based modeling approach can provide mechanistic insights into how the signal changes across regions and scales (Wu et al. 2018). Physics-based models, although unbiased towards out-of-sample, tend to have lower precision (Féret et al. 2019, Gumiere et al. 2020). A rapidly growing technique combines both approaches, with so-called physics-informed machine-learning models showing improved prediction accuracy and precision in situations with low data coverage (Willard et al., 2022). However, a thorough mechanistic understanding of the liana signal spanning multiple scales is a prerequisite for applying such hybrid classifiers.

We explore the global liana spectral signal across leaf, canopy, and stand scales in the visible, near-infrared, and shortwave-infrared spectrum (400 - 2500 nm). Our goal is to understand the mechanisms behind the spectral signal of liana leaves and canopies, compared to host trees, by using radiative transfer models to link species traits to reflectance. Our objectives include: 1) determining if a consistent liana spectral signal exists at various scales using global datasets and different remote sensing tools; 2) identifying traits that create a liana signal via radiative transfer modelling; 3) validating results with independent field data; 4) quantifying trait importance; and 5) evaluating the effectiveness of spectral indices and classifiers in detecting lianas at different scales using simulated spectra.

## Methods

### Overview

To test all objectives, we compiled liana reflectance data in the 400 - 2500 nm range from multiple sites, sensors, and scales (Figure 1, Table S1). We first compared liana reflectance at the leaf level with other plant growth forms and examined case studies of successful discrimination between tropical lianas and host trees on canopy and stand scales (objective 1). Next, we inversely fitted mechanistic radiative transfer models to quantify differences in leaf biophysical, leaf chemical, and canopy architectural properties between lianas and host trees and validate them with ground measurements (objectives 2 & 3). We assessed the relative importance of each trait in generating the liana spectral distribution through iterative model experiments (objective 4). Finally, using forward simulations of the fitted models, we evaluated the potential of different platforms to discriminate lianas and trees under various scenarios (objective 5).

**Figure 1.**
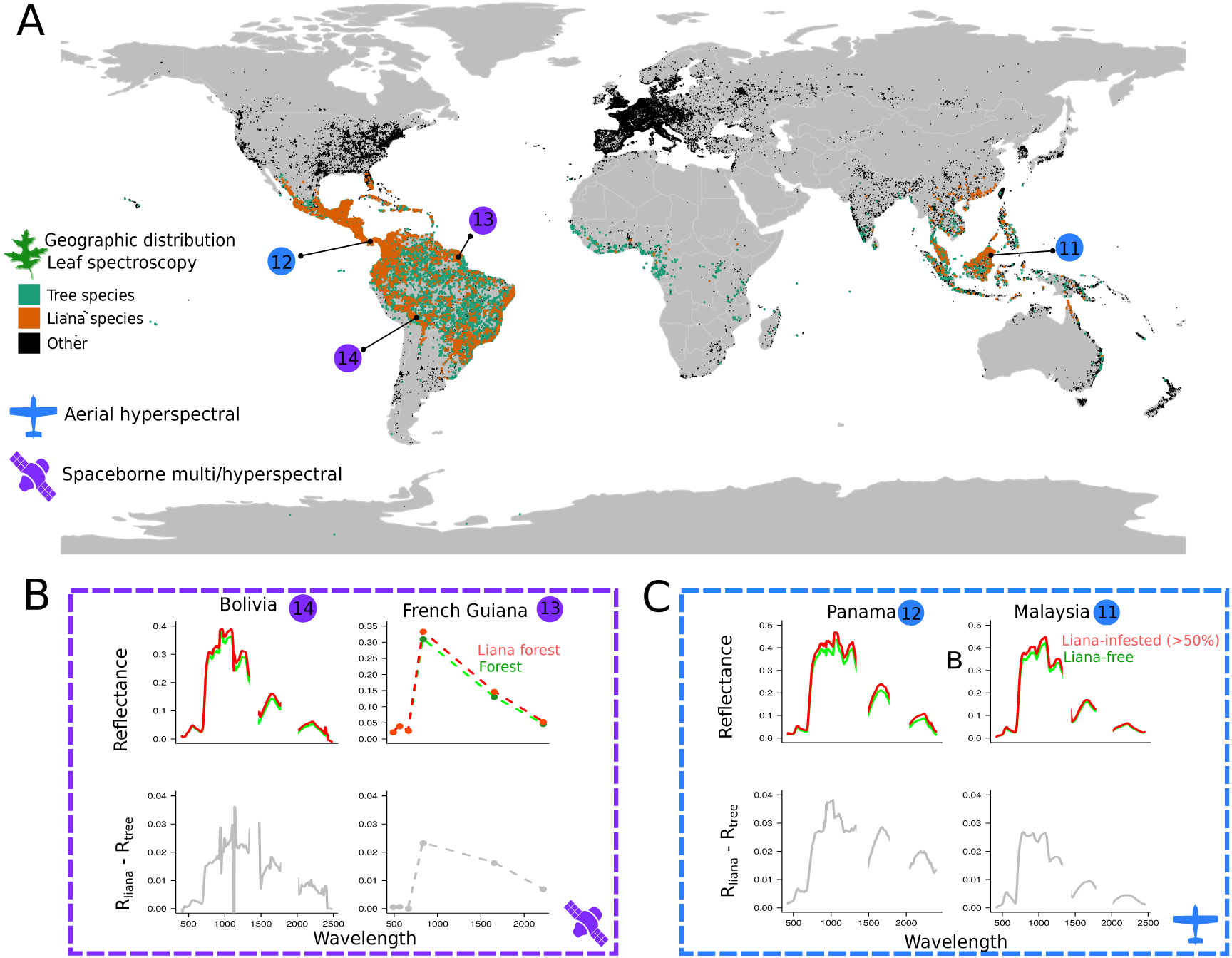
Datasets used in the study to compare the tropical liana and tree spectral signals. Panel A shows the geographic distribution of plants species in the leaf spectroscopy dataset, including the focal tropical liana (orange) and tree species (green). Note that no sampling was conducted in Africa, but three species are common to both the Americas and Africa: *Parinari excelsa, Symphonia globulifera* and the liana *Byttneria catalpifolia*. The locations and remote sensing platform types of canopy scale data are given in blue and purple with reference numbers corresponding to Table S1. Panels B and C display liana spectral signals from airborne and spaceborne sensors at focal sites. Top rows show surface reflectance signals of lightly infested forests (R_tree_) and heavily-infested forests (R_liana_) at each location. Bottom rows show average differences between R_lianas_ and R_trees_.

This paper integrates diverse data from 17 independent sources spanning multiple sites and scales. Due to space and legibility concerns, details on data, field, and laboratory protocols are provided in the supplemental information (SI) and the original published sources.

### Study sites, species and data

#### Spectral datasets

The spectral data (Figure 1, Table S1) span three distinct spatial scales.

*Leaf scale (<<1m^2^ scale)* data encompassed 11 published reflectance spectra datasets covering 5424 leaves of 720 species. Our focus is primarily on tropical species, with 648 tropical tree species and 66 liana species. To place liana leaf reflectance globally, we also included data on 72 crops, shrubs, and herbaceous species in the SI, as all groups could potentially lead to confounding in large classification studies (e.g., Tropek et al., 2014). In all datasets, reflectance was measured in the 300 – 2500 nm range (<2 nm step) with laboratory or field spectroradiometers (Table S1 and Appendix S1). Unique species counts were obtained by cross-referencing names with the GBIF database and tallying unique GBIF keys. Geographic distributions, were also sourced from GBIF, and cleaned following Zizka et al. (2019).

*Canopy scale (1-4 m^2^ scale)* data were obtained from two published airborne hyperspectral campaigns in Panama and Malaysia, covering reflectance from 999 tree crowns with varying liana infestation levels. Further details on the crown scale data, sites and instruments can be found in Appendix S1, Chandler et al. (2021) and Marvin et al. (2016).

*Stand scale* (900 m^2^ scale) data used satellite imagery from previous studies to contrast liana-infested and liana-sparse forest patches in Bolivia and French Guiana (Foster et al., 2008; Tymen et al., 2016). In both cases, we obtained the original images used, which included a 30-m resolution Landsat Thematic Mapper (L1TP) image (French Guiana) and hyperspectral Hyperion imagery from NASA’s EO-1 satellite (Bolivia). Reflectance data span over 775 hectares of forest (40.5 in French Guiana, 734.76 in Bolivia) and was delineated in low liana density forest and adjacent heavily infested stands (Foster et al., 2008; Tymen et al., 2016). In French Guiana, the liana forest was delineated using lidar, aerial photography, and field campaigns, while in Bolivia, it was contrasted with random forest patches using high-resolution videography and field surveys. Both images were georeferenced and surface reflectance corrected. The Hyperion image was corrected for the smile effect. Detailed information on the methods and sites can be found in Tymen et al. (2016), Foster et al. (2008), and Appendix S1.

### Leaf and canopy traits

To achieve objectives 2 and 3, we combined published and newly collected field data on leaf and canopy traits, including data on lianas and host tree species from Panama and Malaysia.

#### Leaf biochemical and biophysical trait data

We obtained measurements of leaf Chlorophyll (C_ab_, μg/cm^2^), Carotenoids (C_ar_, μg/cm^2^), water content (C_w_, g/cm^2^), leaf thickness (μm) and leaf mass area (C_m_, g/cm^2^). Trait measurements followed typical protocols where pigment contents were determined from leaf discs punched from fresh leaves, and water content was measured by recording fresh weight before drying. A subset of the data included matched leaf spectral reflectance and physically measured traits for the same leaves (Table 1). The paired leaf data were used to benchmark the inversion of leaf radiative transfer models (RTMs; see appendices). Average trait values for lianas and trees were also available for dataset 3 in Sánchez-Azofeifa et al. (2009a). Finer details on sampling, extraction, and quantification can be found in Appendix S1.

**Table 1:**
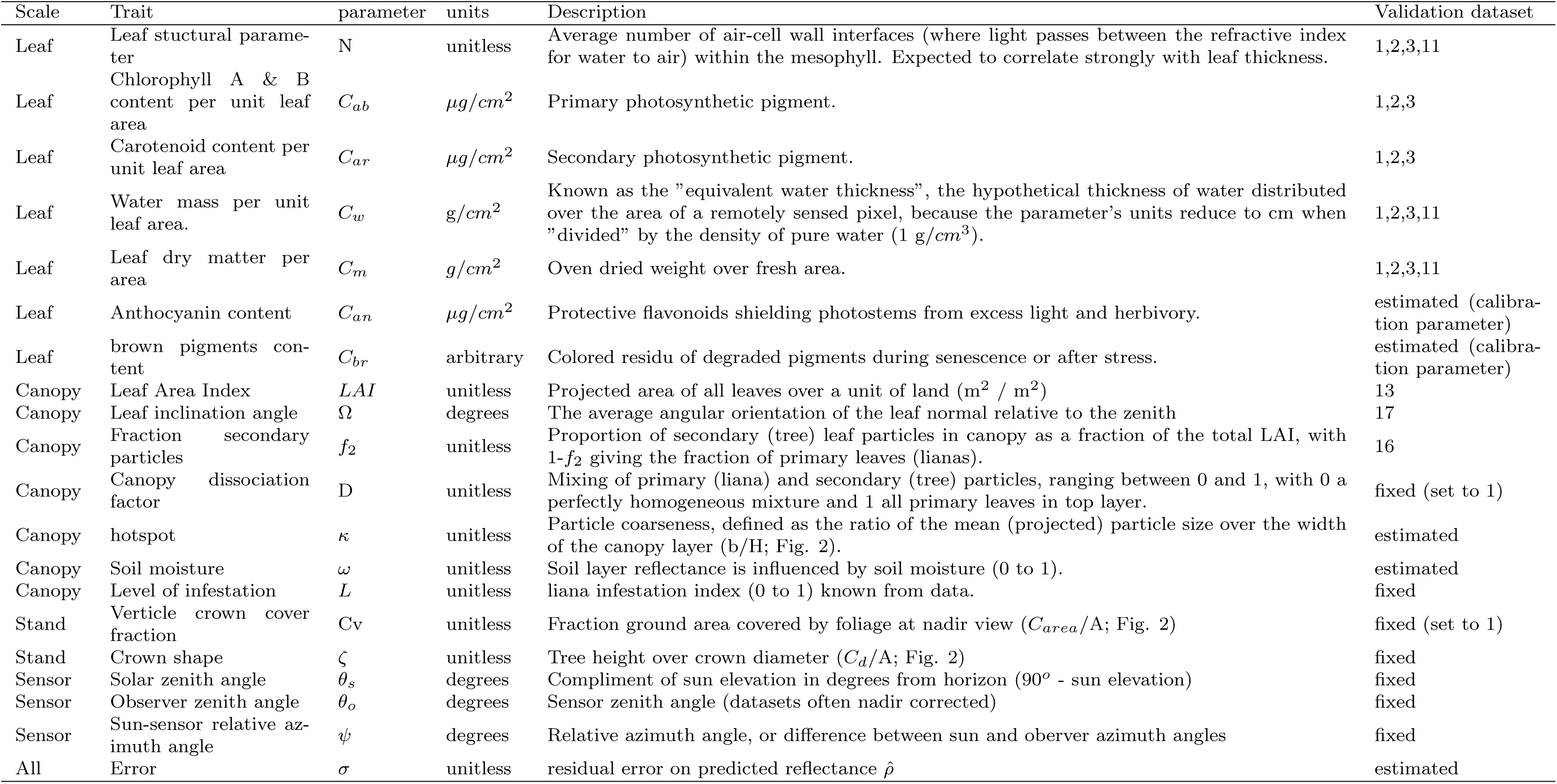
Model parameters and associated canopy and leaf traits.

#### Plant area index (PAI)

The contribution of liana leaves to PAI in infested canopies was estimated using a plant canopy analyzer (LAI2000, LI-COR) (dataset 16 in Appendix S1). The estimation capitalized on a liana removal experiment in central Panama, where the contribution of lianas to PAI was estimated from the difference between control and removal plots (Appendix 1). Details regarding the LI-COR measurements are given in Appendix S1 and Rodríguez-Ronderos et al. (2016), while van der Heijden et al. (2015) describe the removal experiment. Note that we use PAI to refer to empirical measurements, and LAI to refer to the PROSAIL model parameter as per convention.

#### Leaf angles

Leaf angle distribution was measured for canopies of 40 tree species and 10 liana species in Panama (dataset 17). Samples were taken in the canopies of trees and lianas using leveled-digital photography (Ryu et al. 2010, Pisek et al. 2011). Images were taken at different locations, including canopy cranes, telecommunication towers, eddy-covariance flux towers, and canopy gaps. We measured sun and shaded leaves for trees, but only sun leaves for lianas, as the vast majority of liana leaves are in the canopy (Avalos et al. 1999, Medina-Vega et al. 2021a). Leaf angles were defined as the angular orientation of the leaf surface normal relative to the zenith (Ross 1981) and measured from digital pictures using imageJ (Rueden et al. 2017). Whenever the leaves were compound, the angle from the petiole to the furthest leaf tip relative to the zenith was taken as the leaf angle. A total of 1540 individual leaves (646 tree and 894 liana leaves) were measured.

### Radiative transfer models (RTMs)

To address objectives 1-5, we utilized RTMs from the PROSPECT and SAIL families, which have been extensively validated and applied in reflectance modeling studies (e.g. Zhang et al. 2005, Shiklomanov et al. 2016, Wu et al. 2018). All models were programmed in R, optimized for speed following Visser et al. (2015), and refactored in C++ using Rcpp (Eddelbuettel and François 2011). Model code is available as the open-source R-package ccrtm (coupled chain radiative transfer models; version 0.1.6; Visser, 2020). We provide a short overview as these models are exhaustively described elsewhere (see e.g. Jacquemoud and Ustin, 2019).

#### Leaf model

Within the PROSPECT family of models, we evaluated PROSPECT 5, 5b, and D (Féret et al. 2017). for their ability to capture independently measured trait data. We fitted each model to reflectance data for individual leaves and selected the model (5, 5b, or D) that closely reproduced the validation data. Traits, including water mass per unit area, leaf mass area, chlorophyll content, and carotenoids, were validated using paired reflectance and trait data from datasets 1, 2, and 11. Leaf thickness (µm) indirectly validated the leaf structure parameter (N) in datasets 1, 2, 3, and 11. Some parameters, such as brown pigments (C_br_) and anthocyanin content (C_an_), lacked validation data and were treated as calibration parameters: they were only kept if their inclusion improved both the quality of fit and the validation of other parameters (see also Féret et al., 2017). Details on the fit procedure are given below, and all parameters and their units are given in Table 1. Our benchmarks among leaf models showed PROSPECTD performed the best with regards to reducing errors in both the fit and validation datasets (Appendix S4 & 6) and therefore further analyses focus solely on PROSPECTD inversions.

#### Canopy model

We adapted 4SAIL2, which can represent vertically separated vegetation types while maintaining analytical tractability (Verhoef and Bach 2003, 2007, Zhang et al. 2005). Lianas typically form a canopy on top of their host crown (Stevens 1987, Avalos et al. 1999), with empirical studies finding the vast majority of liana leaves in the upper-most leaf layers (Medina-Vega et al. 2021b). This vertical mixing of vegetation types has nonlinear impacts on reflectance, rendering linear mixing inappropriate (Asner 1998). The 4SAIL2 model simulates the reflectance of two-component canopies with distinct vertically separated layers, accounting for differences in leaf optics, optical thickness (LAI), and leaf inclination angles (Braswell et al. 1996). We provide a short description specific to the liana-host tree model implemented here in Figure 2.

**Figure 2.**
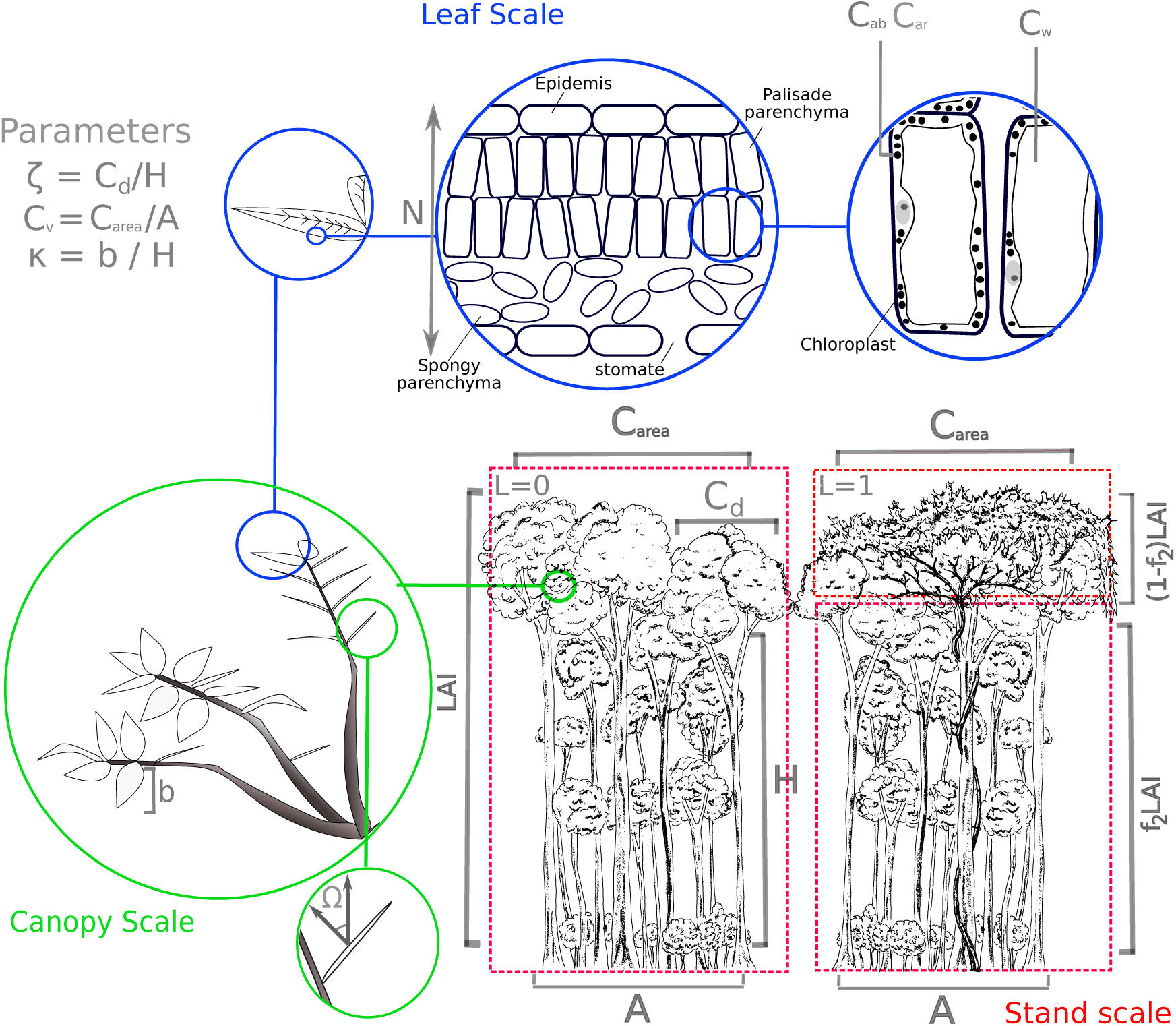
Illustration of the coupled radiative transfer model, merging PROSPECT and FOURSAIL2 to study light behavior in vegetation canopies. PROSPECT addresses leaf radiative transfer, using semi-transparent stacked plates to represent leaf absorption (α), transmittance (τ), and reflectance (ρ). It employs the generalized plate model to determine wavelength-dependent transmissivity (k(λ)), based on empirical absorption coefficients and leaf biochemical concentrations (see table 1). FOURSAIL2 simulates bi-directional reflectance at the canopy level, using differential equations for direct and hemispherical radiance. It splits the canopy into two vertical and infinite horizontal layers with randomly distributed leaves. The model integrates leaf properties (α, ρ, τ) from PROSPECT and allows for different particles, either layered or mixed, using the canopy dislocation parameter (D). In our study, D was set to 1, placing liana leaves predominantly above host leaves. Canopy layers are defined by sensor-sun geometry (θ_s_, θ_o_, ψ), affecting radiance flux length and the angular projection of leaf area. This projection relies on mean leaf angle (Ω_1_, Ω_2_) and leaf area index (LAI), split between top (liana; 1-f_2_) and bottom (tree; f_2_) layers. The model includes a soil layer with variable reflectance and a moisture parameter (ω), and considers the hotspot effect, where equal observer and solar zenith angles increase reflectance. The hotspot’s intensity is calculated as canopy coarseness (leaf width to canopy height ratio, κ = b/H), and the model also accounts for crown cover (Cv) and shape (ζ). However, we only study dense tropical forests, so Cv was set to 1, simplifying the model by negating ζ’s impact on reflectance. Illustrations by M. D. Visser.

#### Coupled chain models

the PROSPECT and SAIL families of models can be coupled together in a chain, typically known as PROSAIL, where the outputs from the former are used as inputs for the latter. Here, we coupled PROSPECT with 4SAIL2, to simulate canopy and stand-scale reflectance in liana-infested forests. The full PROSAIL2 model, in its most complete form, is given by:

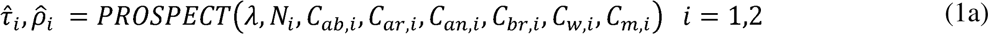

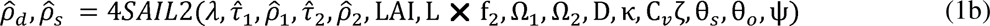

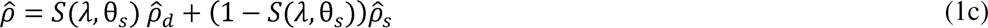

here 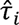 and 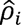 are the predicted leaf transmittance and reflectance (*i* = 1 for lianas, 2 for trees) at wavelength λ, while 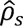 and 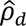 represent bidirectional (direct) and hemispherical directional reflectance (diffuse). The parameter f_2_ and *L* divide the total leaf area (LAI) between the top and bottom layers, with *L* being the infestation intensity index (ranging between 0 and 1). The optical thickness of the top liana layer is assumed to be proportional to the liana infestation intensity (*L* × (1 − f_2_)). This results in a maximum optical thickness of (1 − f_2_) for the top layer when *L* = 1, and the canopy reverts to a single layer with only tree leaves when the infestation is zero (*L* = 0). The fraction of LAI assigned to lianas is then LAI × L (1 − f_2_). The final predicted reflectance of the target pixel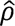 depends on the fraction of diffuse light, S(λ, θ_s_), at wavelength λ and solar zenith angle (θ_s_) (following Danner et al., 2019; François et al., 2002). Parameters detail and units are given in Table 1 and Figure 2.

### Inverse modelling of the liana signal

Spectral inverse modeling aims to estimate physical parameter values consistent with observed reflectance data (objective 2). We employed a Bayesian framework for spectral inverse modeling (Shiklomanov et al. 2016), where coupled chain RTM-models are particularly useful, as models fitted at one scale can inform models at another scale. We first fit the PROSPECTD model to all 5424 individual leaf-reflectance spectroscopy data, decomposing variation among groupings, using a hierarchical modelling approach:

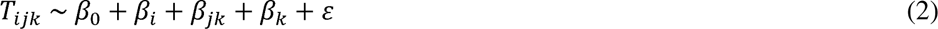

where *T_ijk_* represents one of the RTM model parameters in equation 1a, fit for growth form *i* (liana or tree) from species *j* nested in study *k*. We included study to account for variation due to factors associated with specific methodologies, such as the usage of different spectroradiometers (Meireles et al. 2020). All β parameters were assumed to be normally distributed with uninformative hyperpriors. We utilized leaf-level means and variances to inform priors for leaf parameters at canopy and stand scales, assuming them to be normally distributed. The informative priors for both lianas and trees largely overlapped, with their means falling within one standard deviation of each other in every instance. All other parameters had uniform flat priors (see Appendix S2 for details). For models fit at all scales, we sampled the joint posterior distribution of each RTM with Monte-Carlo Markov Chain (MCMC) methods using the differential evolution adaptive metropolis algorithm, implemented in the ‘BayesianTools’ R package (Hartig et al. 2019). In the inversion procedure we excluded the multispectral data from French Guiana (dataset 13) because parameter uncertainty is sensitive to spectral resolution (Shiklomanov et al. 2016). Therefore, we focus on hyperspectral data (datasets 12, 13, and 15), and included each sensor’s spectral response functions in the fitting procedure. Finer details on model inversion are given in Appendix S2. We conducted several model robustness checks, including simulations to determine parameter identifiability and the effect of measurement error (Appendix S3).

### Spectral differences and the detection of the liana signal

Kullback–Leibler divergence (KLD) is a measure from information theory that quantifies the difference between probability distributions, commonly used for optimization in machine learning (Kosheleva and Kreinovich 2017). As a measure of feature discernibility for classifiers, KLD helps explore differences in spectral distribution between plant groups, the impact of traits on spectral distributions, and the theoretical ability to distinguish lianas from trees (objectives 1, 4, 5). At the leaf level, we used KLD to compare different plant groups, with tropical liana leaves as the reference (0). KLD is interpreted as the information lost when encoding the liana spectral distribution (0) with another plant group’s distribution (k). KLD was defined as:

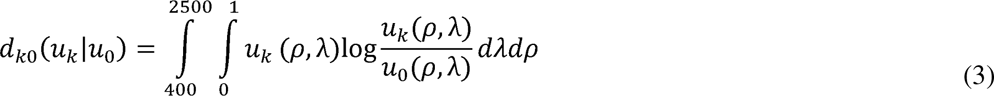

where *u*_k_(ρ,λ) is the probability of observing a reflectance value ρ at wavelength λ, and *u*_0_(ρ,λ) is a reference distribution. Assuming normality, the spectral distribution can be estimated with the wavelength specific mean and standard deviation, *μ*(*λ*) and *σ*(*λ*) respectively. In this case KLD (eqn. 3) reduces to:

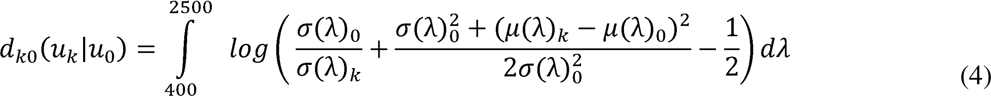

The smaller the KLD value, the more comparable the groups are, decreasing the likelihood of robust classification (Kosheleva and Kreinovich 2017). The expected KLD between two identically distributed signals with equal means and variances is, by definition, zero.

#### Model sensitivity experiment

KLD quantifies the loss or gain of information due to individual traits by comparing modelled spectral distributions that include or exclude one or more parameters. This provides a metric of individual traits’ importance (objective 4). KLD was calculated with predicted reflectance 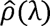 as the mean and variance estimated from the residual error from the inverse fits (σ, Appendix S2). Starting with a model parameterized with mean tree leaf and canopy traits as the focal distribution 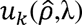, we iteratively changed single parameters to liana-specific values and calculated the change in KLD using the liana spectral distribution as the reference distribution 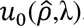. This determines the parameters with the greatest information loss or gain towards the liana spectral distribution. To understand influential parameters at different wavelengths, we also evaluated the KLD at each wavelength separately and assessed combinations of two parameters to identify interactions that most strongly recreated the liana pattern.

#### The theoretical ability of classifiers to detect the liana signal

In theory, sensors and spectral indices that sample at bands maximizing KLD between trees and lianas optimize discernibility between these groups. Hence, sensor sampling regions of maximum KLD are expected to provide optimal information for modeling and classification (Wang et al. 2015). Therefore, we compare how well the spectral response functions of several common multispectral remote sensing platforms maximize KLD. Given the strong spatial clustering of lianas at different scales (Ledo and Schnitzer 2014) and their varying infestation patterns from a) incidental crown infestation (Marvin et al. 2016) to b) arrested succession (Schnitzer et al. 2000) and c) liana forests (Tymen et al. 2016), we investigate how the spatial aggregation scale interacts with sensor spatial resolution in determining the expected KLD. We simulated 1000×1000 pixel hyperspectral image scenes with various scales and autocorrelation degrees in liana infestation for scenarios (a-c). We started with the finest scale (1 m^2^) as pure signals and then recalculated the expected KLD between feature (liana) and non-feature (liana-free) areas, gradually reducing spatial resolution from 1×1 m to 250×250 m via resampling, repeating this 50 times per scenario. Each scenario maintained a roughly equal number of feature pixels. The sensors modeled included Landsat TM, Worldview 2’s WV110 camera, and Hyperion.

Note that we focus solely on the aspects of sensor choice readily controllable by the user. We explicitly ignore atmospheric effects that decrease the signal-to-noise ratio of most sensors (Landgrebe and Makaret 1986). Consequently, the results should be considered a “best-case scenario”.

## Results

### Objective 1: Does a globally consistent liana signal exist at the leaf, canopy and stand scales?

The distribution of liana leaf-scale reflectance, measured from 718 leaves from 66 species, exhibited significant overlap with trees, shrubs, crops, and grasses (Figure S6). This finding was further supported by inversely fit PROSPECT leaf models (Figure 3 A-C). Kullback-Leibler divergence (KLD) analyses on spectral distributions and predicted posterior distributions confirmed the indistinguishable nature of tree and liana leaf reflectance across all wavelengths (Figure 3). In contrast to leaf reflectance, however, predicted posterior distributions of leaf transmission and absorption demonstrated substantial differences, particularly in the NIR and SWIR regions (Figure 3 E-J). Liana leaves had higher transmission and lower absorption throughout the evaluated spectrum, leading to pronounced distributional divergence (Figure 3 G & J).

**Figure 3.**
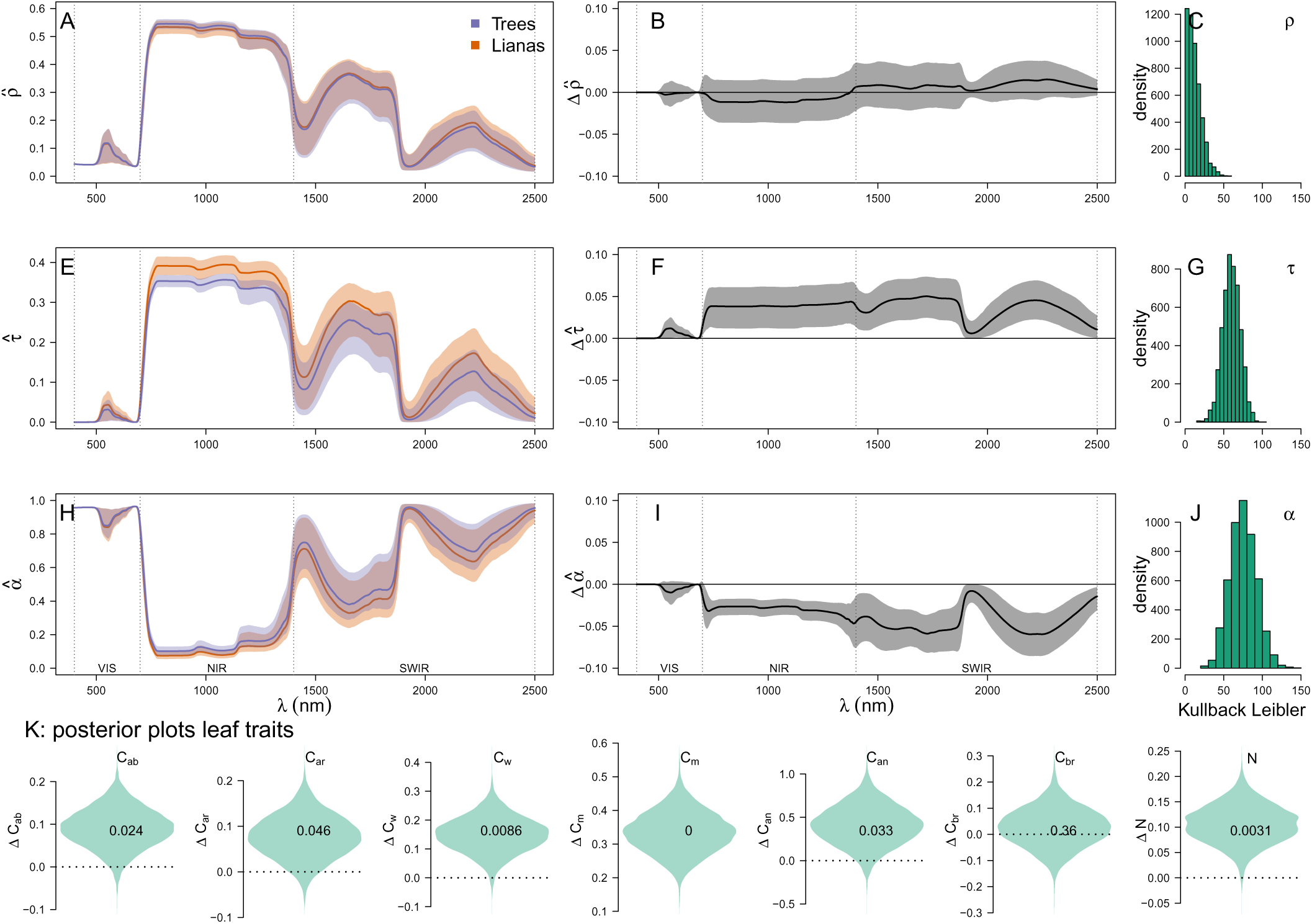
Spectal properties of liana and tree leaves in the solar spectrum (400 to 2500 nm). Panels A to J show the predicted posterior distributions of reflectance 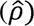, transmission 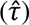 and absorption 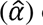 of liana and tree leaves (A, E, H), and the difference between the posterior distributions (B, F, I) – here solid lines depict means and the shaded envelop shows 95% credible intervals. Kullback-Leiber Divergence (KLD) quantifies how the overall spectral probability distribution of trees diverges from liana for reflectance (C), transmission (G), and absorption (J) – withe zero indicating an identical distribution. The violin plots in panel K, show differences between lianas and trees in predicted posterior distributions of leaf traits. From right to leaf; chlorophyll content (C_ab_), carotenoids (C_ar_), water mass per unit area (C_w_), leaf mass area (C_m_), anthocyanin content (C_an_), brown pigments (C_br_) and leaf structure (N). Fractions within each violin plot incate the fraction of posterior samples that overlap with zero (i.e. no difference).

Liana-infested canopies showed consistently higher reflectance than liana-free canopies at both canopy and stand scales across various sites, platforms, and wavelengths (Figure 1B & C). The reflectance difference (**R_lianas_** − **R_trees_**) had an average 89% correlation between sites (range 74% to 99%) when matched to Landsat 5TM bands (used in French Guiana). For hyperspectral images at overlapping wavelengths, the mean correlation for reflectance difference was 96% (range 95%-98%). These results reveal a distinct and detectable spectral signal for liana reflectance at canopy and stand scales, stable across instrumental noise and sensor differences.

### Objective 2: Inverse modelling of the liana spectral signal at the leaf, canopy and stand scales

The leaf signal was accurately reproduced by PROSPECTD exhibiting a low overall error (R^2^ > 0.94 for 5424 individual leaves; Appendix S4). Inverse fits of the PROSAIL2 model to canopy and stand scale data closely approximated the spectral reflectance of heavily infested (>50% cover) and lightly infested canopies (<50%) in Bolivia, Malaysia, and Panama (R^2^ > 0.98; Figure 4A-C). The model also successfully replicated the difference in reflectance between heavily and lightly infested canopies (**R**_lianas_ − **R**_trees_), explaining between 46% and 92% of the variation (inset plots in Figure 4A-C). Model posteriors can be found in Appendix S5.

**Figure 4.**
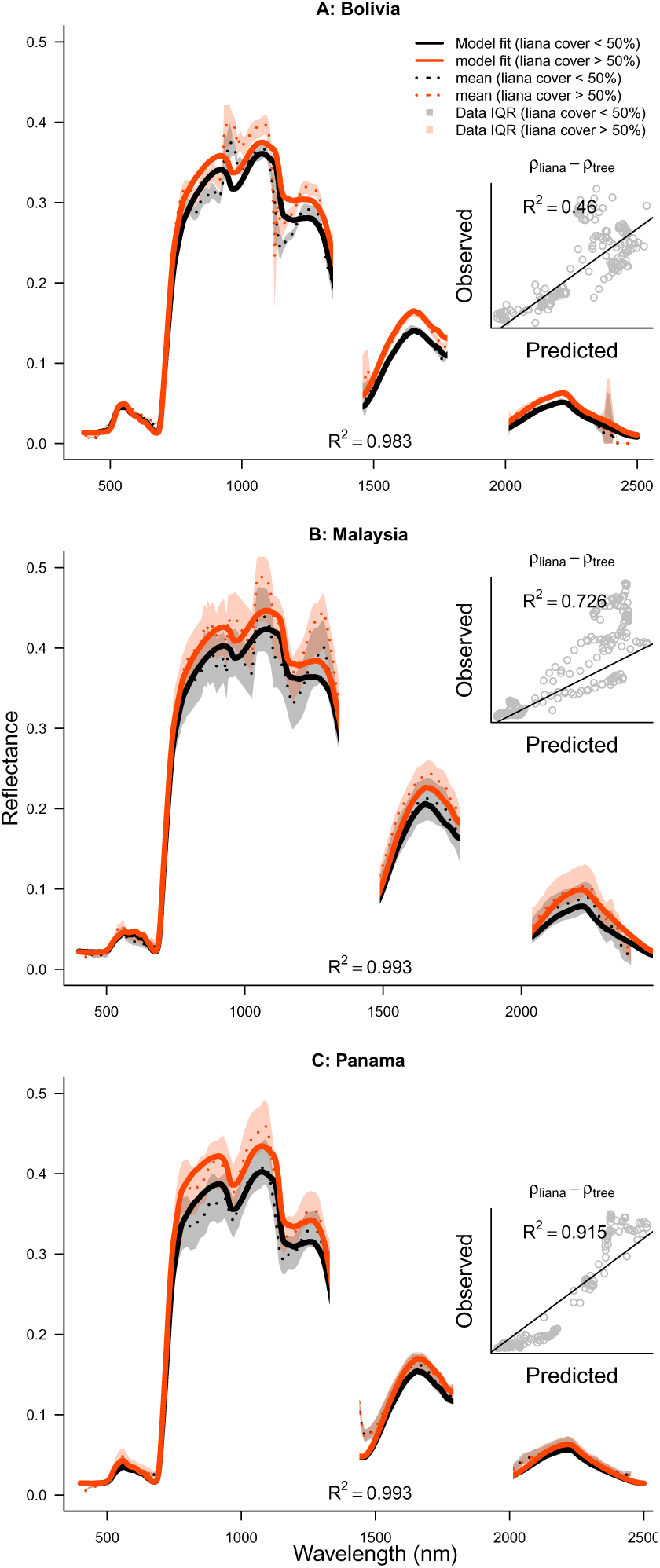
Model predictions for high and low levels of lianas coverage compared to their observed reflectance from airborne and satellite hyperspectral signals. Dotted lines and colored envelopes give mean and inter quartile range (IQR) respectively. Model fits performance was uniformly high, as measured by the coefficient of determination (R^2^ > 0.98). Inset plots show model predictions of the differences in the signal between high and low liana coverage, which all compared well with the observed difference (R^2^ **=** 0.46, 0.73, 0.92), respectively for Bolivia, Malaysia and Panama).

### Objective 3: Model validation

The PROSPECTD model, fitted to leaf reflectance data, accurately and unbiasedly replicated independently measured leaf traits such as C_ar_, C_ab_, C_m_, and C_w_, with observed and predicted traits showing agreement ranging from R^2^= 0.4 to 0.73 (Appendix S6). The leaf structural parameter strongly correlated with leaf thickness (R^2^= 0.54), suggesting it captured meaningful physical variation.

Figure 5 depicts the predicted differences between lianas and trees based on inverse fits of PROSAIL2 (black dots and whiskers) for parameters showing non-overlapping credible intervals (CI) at least once across sites. The pattern of differences between lianas and trees in inversely estimated parameters was generally consistent across sites, although not all parameters exhibited significant differences at all sites based on 95% CI (Figure 5). Regarding leaf biophysical and chemical parameters, all models indicated that lianas had more cheaply constructed leaves on average, characterized by thinner leaves, lower photosynthetic pigment content (Chlorophyll and Carotenoids) per unit leaf area, and lower dry mass and water mass per unit leaf area. Note that water mass C_w_ (g/cm²) should not be equated with leaf water content (%): lianas had higher water content compared to trees (77% vs 70% on average). Inverse fits with PROSPECTD confirmed the independent predictions of PROSAIL2 whenever leaf-level spectral data were available (blue dots and whiskers in Figure 5), except for C_m_ in Malaysia, which was lower according to PROSAIL2 compared to C_m_ estimated from leaf reflectance (PROSPECTD) and direct measurements. However, all estimates of C_m_ in Malaysia aligned with the general trend of lower leaf mass per unit area for lianas compared to trees (Figure 5). At the canopy and stand scales, the models predicted that trees contribute 66% (f_2_) to the total LAI across sites. Indicating that liana leaves contributed an average of 34% (1-f_2_) when a tree crown is fully infested, with site averages ranging from 16% to 55% (Figure 5). Liana leaves were also predicted to have leaf angles that were 23.5% flatter compared to trees on average, with parameter Ω_1_ being 4% to 59% lower than Ω_2_ across sites (Figure 5).

**Figure 5.**
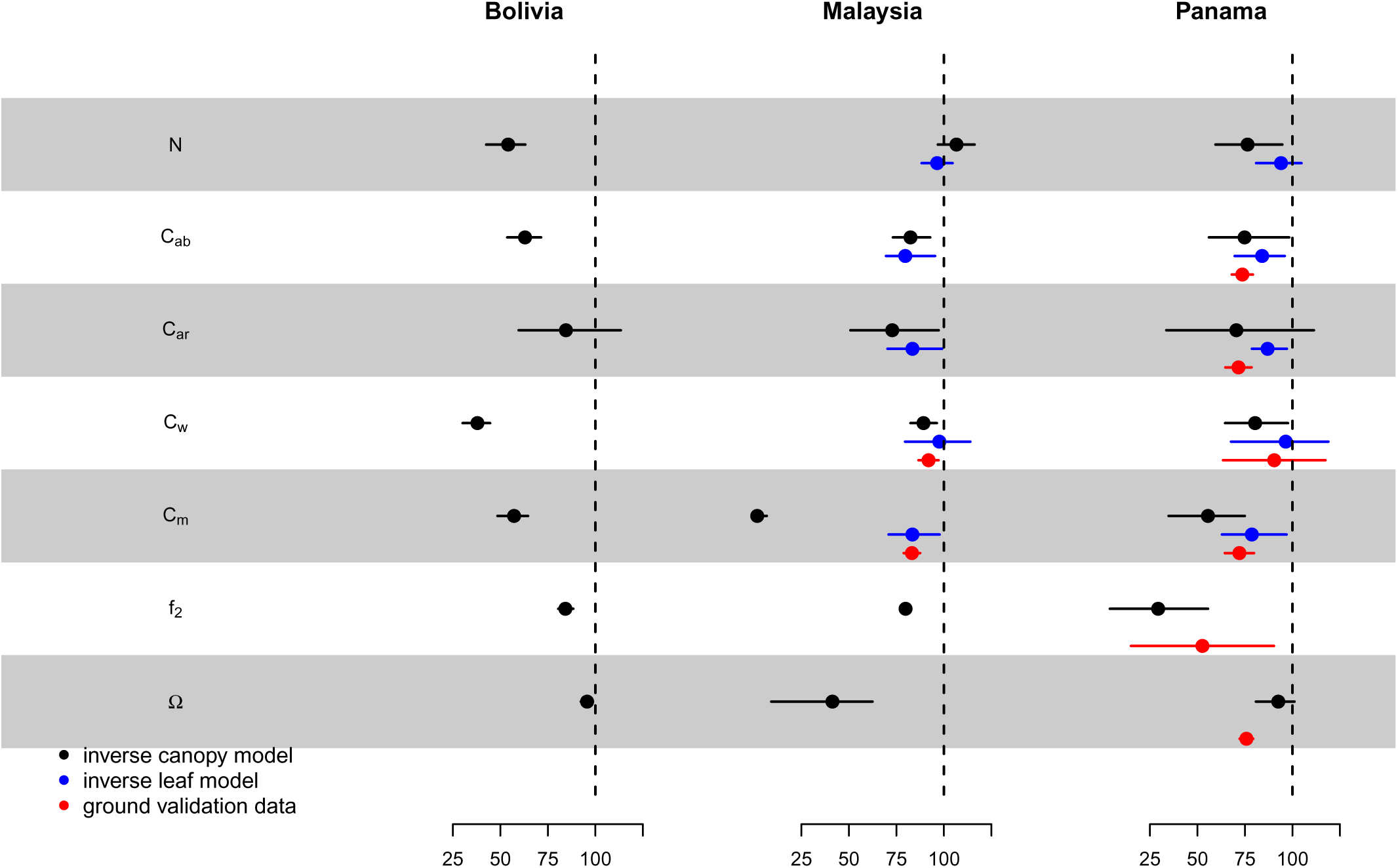
Leaf and canopy traits of liana-covered and liana-free forest canopies. Inverse estimates for lianas and trees from full canopy models are compared at each site (black) with independent leaf model data from spectrometers (blue) and, where available, independent field and lab measurements (red). Traits are shown as percentages of average tree traits for easy comparison, irrespective of units. Absolute values and ground truth comparisons are in Appendix S6. The fraction of leaf area index due to lianas is noted as 1-f2. Mean values and 95% credible intervals are represented with dots and whiskers. Parameters not significantly different between groups across any sites are omitted. Full details on parameters and their posterior distributions are in Appendix S5.

The inverse fits with the PROSAIL2 model were supported by independently collected verification data where available. Lab measurements of leaf thickness, Chlorophyll and Carotenoids content, dry mass, and water content per unit area for liana leaves were all found to be lower on average for liana leaves (red dots and whiskers in Figure 5). Canopy and stand scale validation data also showed consistency with the model predictions. The estimation of the fraction of PAI attributed to trees vs lianas (f_2_ vs1-f_2_) using LI-COR data in plots where lianas were removed showed that trees contribute 65.3% (f_2_ = 0.653 ± 0.087 SE) to the PAI when their canopies are infested. Indicating that lianas contribute on average 34.7% - a value close to the inverse model estimate. Additionally, the inverse model results predicted flatter leaf angles for all sites (Figure 5), and measurements conducted in Panama confirmed that lianas have, on average, 26.2% flatter leaf angles compared to trees (mean angle of 27.9° ± 0.76 SE vs 37.8° ± 0.64 SE).

### Objective 4: what causes the liana signal?

The influence of parameters on the liana spectral distribution varied depending on the solar spectrum region (Figure 6A). In the visible spectrum, leaf angles (Ω) and photosynthetic pigment content (C_ar_, C_ab_) played significant roles, with flatter leaf angles and lower pigment content leading to increased reflectance. In the near-infrared, leaf angles (Ω), leaf mass area (C_m_), and leaf structure (N) consistently showed importance, although the relative significance of leaf angles (Ω) versus leaf mass area (C_m_) differed among sites (Figure S8A, C, E). Leaf water content (C_w_) became increasingly influential with wavelength and dominated the signal in the short-wave infrared (Figure 6A), particularly at the tropical dry site in Bolivia (Figure S8A). The most influential traits overall were leaf mass area (C_m_), leaf angles (Ω), and leaf water content (C_w_) (Figure6B). Leaf angles (Ω) were consistently important across sites, while the importance of C_m_ and C_w_ varied among sites (Figure S8B, D, F). Substituting the leaf structural parameter (N) of trees with that of lianas resulted in predictions further deviating from the liana spectral distribution (negative contribution), especially when interacting with other parameters (light colors in Figure 6B).

**Figure 6.**
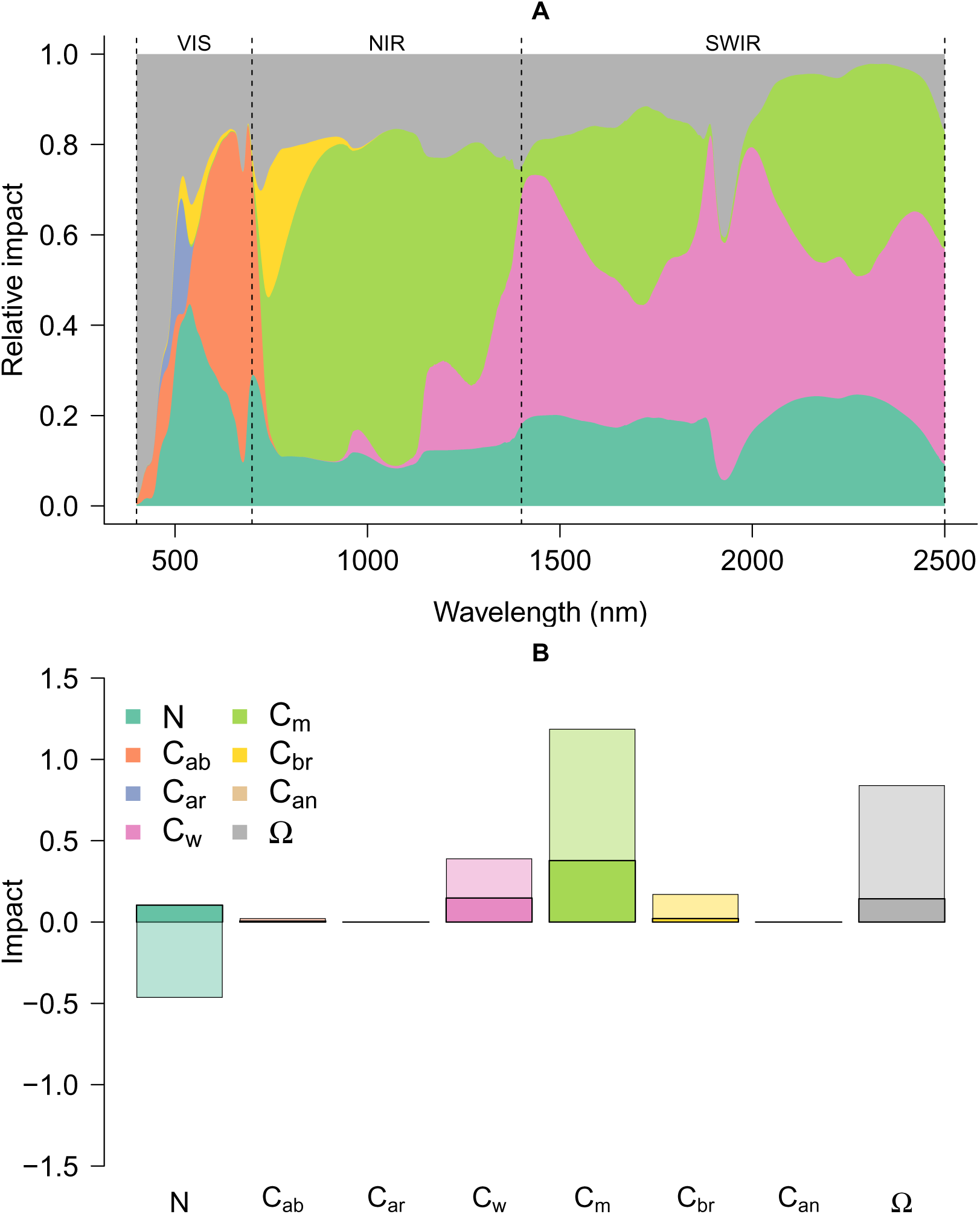
The relative importance of different traits in generating the liana signal across sites. Here, spectral traits of the top layer were initially set as tree traits and gradually replaced with liana traits either one by one (additive) or in pairs (interactive effects). The Kullback-Leiber divergence score was calculated at each iteration. Panel A displays the average relative absolute impact of changing a single trait in the top layer at each wavelength across all sites. Panel B shows the total impact of additive (darker colors) and interactive effects (lighter colors). Positive impact indicates a change towards the liana signal, while negative impact indicates a divergence. Figure S7 presents the same data for individual regions (Bolivia, Malaysia, and Panama).

### Objective 5: theoretical limits to the detection of liana

The KLD between liana-infested and liana-free spectral distributions was highest in the NIR and SWIR bands, which are covered by several multispectral spaceborne sensors (Figure 7A). The performance of a sensor in discriminating lianas from trees improved with an increased number of bands within the optimal range (Figure 7B). The spatial resolution of the sensor interacts with liana spatial aggregation (Figure 7C-E). In a crown infestation scenario (liana pixel aggregation ~ 350 m²), the divergence between feature and background decreases rapidly (Figure 7C). For larger spatial aggregation resembling arrested forest succession (2000 m²), the divergence dropped at a similar pace to random feature pixels (Figure 7D). For highly aggregated lianas across extensive forest areas (~30,000 m²), the divergence decreased slower than the random expectation.

**Figure 7.**
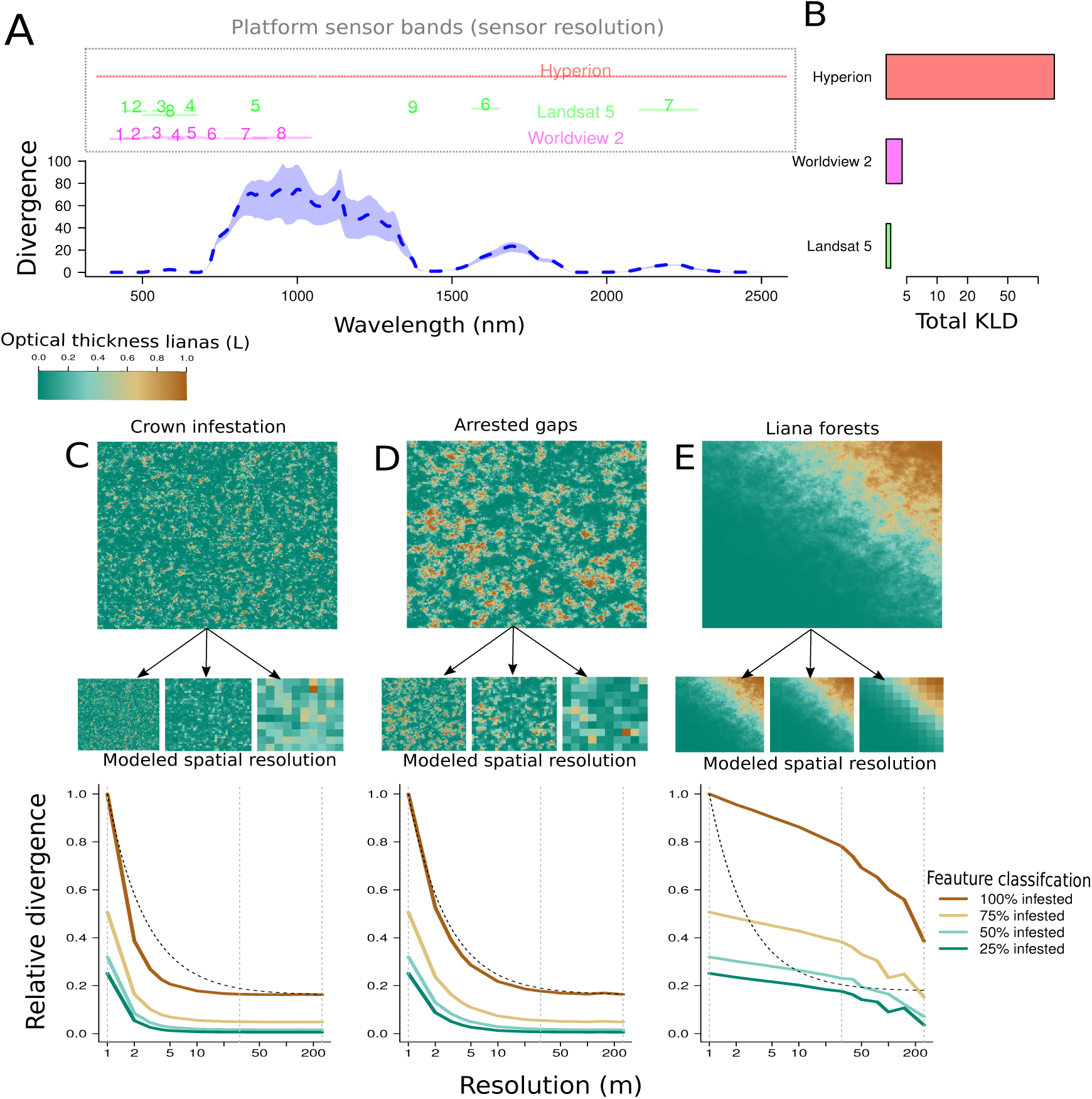
The theoretical capacity of classifiers to distinguish lianas using the expected Kullback-Leiber divergence (KLD). Panel A predicts KLD across wavelengths for 100% liana infestation, with model-based variance estimation, and overlays spectral sampling regions of various remote sensors to assess their effectiveness in liana detection. Panel B presents the cumulative KLD for each sensor, with mean divergences of 0.63 (Hyperion), 0.54 (WorldView 2), and 0.45 (Landsat 5). Panels C-E explore the impact of sensor spatial resolution on liana discernibility at different scales: crown-scale infestation (C), forest gap arrestation (D), and liana forests (E). They include simulated scenes and resampling results, showing the effect on KLD. Liana cluster sizes in these simulations are 350, 2000, and 30,000 m², corresponding to infestation scales from prior studies (Marvin et al. 2016; Chandler et al. 2021; Schnitzer et al., 2000; Foster et al. 2006; Tymen et al. 2016). The colormap indicates liana infestation levels, while line colors in the graphs show different features. Grey vertical lines mark different resolutions, and a black dashed line shows the expected KLD decline for random feature distribution. Each scenario has a roughly equal number of feature pixels.

## Discussion

We demonstrate a unique spectral distribution in liana-infested canopies and forest-stands (Figures 1, 4 & 7), showing higher reflectance than liana-free areas (Figure 1 & 4), particularly in the NIR and SWIR spectrum sections (Figure 7). This distinct spectral distribution, observed across four global sites (Figure 1), has implications for liana evolution and forest energy budgets - and provides a basis for expanded research on lianas at pantropical scales. However, the traits underpinning this distribution might not be exclusive to lianas, posing a clear risk of confusion with other phenomena.

### The mechanisms behind the liana spectral signal

Our analysis reveals a consistent liana spectral distribution across sites, observable at larger scales (≥ 1 m^2^; Figure 1 & 4), but absent in leaf-scale reflectance (<< 1m^2^; Figure 3). Model predictions indicate leaf traits alone do not account for the consistent reflectance trend between liana and tree leaves across sites (Figure 2A & Figure S7). Previous research confirms varying reflectance distributions and optical traits among liana species and sites (Sánchez-Azofeifa et al., 2009a; Asner and Martin, 2012; Medina-Vega et al., 2021a). Therefore, we conclude that leaf reflection alone is unlikely to be critical. In contrast, differences in leaf traits between lianas and trees do affect transmission and absorption profiles (Figure 2) and, by interacting with canopy traits, likely drive the distinct distribution observed across multiple sites and platforms (Figure 1 & 4). Our findings are supported by radiative transfer model inversions at the canopy and stand scales (Figure 5 & 6).

Differences in leaf traits between lianas and trees yield consistent trends in leaf absorption and transmission (Figure 2), and we predict lower leaf absorption and increased transmission for liana leaves at the canopy scale across evaluated sites (Figure S7). Plausible mechanisms for the distinct liana spectral distribution involve reduced light extinction rates in liana-infested canopies. In addition, our model experiment highlights the impact of flatter leaf angles in liana leaves compared to host trees (Figure 5, 6). We propose a two-component mechanism for the “liana signal”: 1) higher projected leaf area, which, at equal leaf reflectance, leads to increased reflected energy at the canopy level, and 2) multiple scattering from greater transmission and lower leaf absorption, which indirectly increases reflectance. The latter is an effect similar to what is hypothesized to cause canopy greening (Wu et al. 2018).

Building on our mechanistic models, we can derive theoretical expectations for the generation of liana canopy-level spectral distribution. The first component arises as lianas retain, on average ~ 24% flatter leaf angles (27.9° vs 37.8°), which should result in roughly an 11% increase in projected leaf area as seen from a polar angle and a proportional increase in the reflected energy. The second “multiple scattering” component arises from liana leaves having lower average absorption (Figure 3 & S7). This results in slower decay of light energy (Appendix S9) and increased scattering, including in the observer’s direction, leading to higher reflectance when integrated across the canopy. Multiple scattering also explains the regions where the liana spectral distribution differs most from trees. Since the projected leaf area does not change with wavelength, its importance is relatively uniform across the spectrum (Figure 6A, parameter Ω). The importance of multiple scattering, however, peaks when leaf transmission is maximized, in the SWIR and NIR parts of the spectrum. This corresponds to the point of maximal discernibility of the liana distribution (Figure 7), and where leaf traits appear the most influential in our sensitivity analysis (Figure 6).

In conclusion, a distinct liana spectral distribution emerges from interactions between multiple leaves within canopies. This multi-leaf dependency explains why the signal is observed at the canopy scale (Figure 1 & 4; Chandler et al 2021, Waite et al 2020, Foster et al 2008, Tymen et al 2016) and the lack of a consistent signal when only leaf reflectance is considered (Figure 3, Sánchez-Azofeifa et al., 2009a).

### Ecological consequences of the liana spectral signal

By increasing forest canopy albedo (Figure 2), lianas have the potential to impact forest energy budgets and understory light conditions. As tropical forests are typically light-limited and liana abundance is increasing in the Neotropics (Schnitzer and Bongers 2011, Wright et al. 2015), liana proliferation could have broader impacts beyond current knowledge.

Meunier et al. (2021) integrated liana optical properties into the ED2 dynamic vegetation model suggesting several consequences. With approximately 25% liana leaf area, the model predicted an increase in reflected energy of 11.7-17.1% across the PAR and IR spectrum. This resulted in a reduction of 3.6 Wm^−2^ in absorbed energy, leading to darker understories (up to 50% darker) and slightly cooler soils (−0.5°C in the topsoil). These findings highlight the broader impacts of the liana spectral signal on forest functioning, including temperature-dependent processes such as metabolic respiration and photosynthesis. Our results support Meunier et al.’s findings and indicate a global characteristic of higher albedo in liana-infested areas. We believe this shows a direct and urgent need for empirical validation of these model predictions to better understand the potential long-term changes in forested ecosystems with increasing liana prevalence.

### The evolution of a global liana spectral distribution

Lianas consistently display cost-effective leaves with flatter angles than their tree hosts, as shown across diverse tropical regions in Asia and South America (Figure 5). This pattern, uniformly revealed through our inverse modeling at leaf, canopy, and stand scales, is notable given lianas’ independent evolution in various angiosperm families (Putz and Mooney 1991, Gentry 1992). This prompts a fundamental question: why do lianas possess a similar spectral signal across vastly different sites? We propose that this phenomenon may be an example of convergent evolution.

Lianas rely on structural parasitism to access the heights and light conditions required for seed production and dispersal. This, however, increases their host’s mortality (Ingwell et al. 2010, Visser et al. 2018a), which consequently shortens their residency in the canopy. This strongly mirrors pathogen virulence evolution (Anderson and May 1982), and evolutionary theory suggests that lianas should, therefore, evolve to be less harmful to hosts, extending their canopy residency and reproductive time. Such adaptation would involve slower growth and reduced resource usurpation from hosts, entailing minimized host-shading (e.g. Ichihashi and Tateno 2011). Slow growth implies greater investment in leaves (Osnas et al. 2013) and steeper leaf angles for better light penetration, akin to tree strategies that optimize their leaf angles to allow optimal light levels to reach lower leaf layers (Falster and Westoby 2003). The fact that these expectations are not realized, is probably related to the fact that lianas seldom exploit their hosts in isolation: tree canopies are typically infested by two or more liana species (Visser et al. 2018b).

In such multi-parasite systems, theory predicts a “tragedy of the commons”, where any parasite limiting host exploitation becomes susceptible to invasion by a more exploitative variant (Frank 1996). This leads to heavy competition for resources and time at the expense of the host: reproductive success requires maximized light capture and rapid growth to avoid being overshadowed by other lianas. We suggest that this “tragedy of the commons” is a common evolutionary force among forest lianas, driving rapid growth, turnover and metabolism and resulting in lower leaf longevity and construction costs among sun-exposed lianas compared to trees (Slot et al. 2013, Wyka et al. 2013). In contrast, the most common hosts for lianas are shade-tolerant trees (Visser et al., 2018a), which typically have costlier, long-living leaves and high Leaf Area Index (LAI) values (up to 5 in an individual canopy; Kitajima 1994) - creating a distinct spectral contrast to lianas.

#### Robust detection of lianas

Our simulations highlight two primary insights regarding liana detection. First, many sensor platforms often miss optimal bands distinguishing lianas from trees. Satellite platforms might possess just one optimal band (Figure 7A). Second, as crown infestation levels increase, the liana spectral distribution decays non-linearly (Figure 7C-E). High spatial resolution is essential for detecting lianas within tree crowns, where signal weakening due to spectral mixing occurs faster than expected (Figure 7C). This means that moderate-scale platforms (around 20-30m resolution) are less suitable for detecting crown infestations but should be able to identify larger liana-infested regions where the signal decays slower (Figure 7E). Our simulation results align with previous studies. These two simulation findings echo patterns from earlier studies. Moderate resolution platforms like Hyperion and Landsat effectively identified large liana-dominated areas (Foster et al. 2008, Tymen et al. 2016), whereas high-resolution hyperspectral images were best for detecting heavy tree crown infestations (Marvin et al. 2016, Chandler et al. 2021). Hence, our findings can inform future liana remote sensing research and sensor selection. However, caution is advised as this analysis, being a “best-case scenario”, overlooks atmospheric factors potentially weakening satellite signals (Landgrebe and Makaret 1986).

#### Confounding variables

Our analysis of the liana spectral distribution prompts careful use of classifiers due to potential signal confusion. Notable confounders are:

1. Leaf phenology. Seasonal shifts in phenology can confound classification. Like lianas, younger tree leaves exhibit lower C_m_, C_w_, and pigment concentrations, causing them to resemble liana signals by being more transmissive (Wu et al. 2018).
2. Tree life history. Light-demanding species possess flatter leaf angles and thinner leaves, risking liana misclassification. Concurrently, liana infestation can obscure tree signals, causing mapping inaccuracies (Zhang et al. 2006).
3. Topography and sun-sensor geometry. Another possible confounder is areas sloped towards the sun. These will appear to have relatively higher reflectance across the spectrum.
4. Climate. Some key leaf-trait differences between lianas and trees identified here, lower C_m_, and C_w_, remain but become less pronounced in wetter climates (Sánchez-Azofeifa et al. 2009a, Asner and Martin 2012, Medina-Vega et al. 2021b). As these traits are mostly responsible for the maximal divergent signal from trees in the NIR and SWIR portions (Figure 7), the spectral difference between liana-infested and free canopies may also become less pronounced.

These confounding factors in detecting lianas can be addressed uniquely. Seasonal leaf phenology changes contrast with lianas’ longer-term growth, aiding detection, especially in dry forests where lianas retain leaves longer (Schnitzer 2005). Light-demanding tree species, less prone to liana infestation and with fewer leaf layers (Kitajima 1994, Visser et al. 2018a), can be distinguished using lidar structural data alongside spectral information (sensu Rao et al., 2020). This approach is particularly relevant given the substantial structural changes caused by liana infestation, such as canopy wood area index alteration (Sánchez-Azofeifa et al. 2009b). Furthermore, topography and sun-sensor geometry issues might be mitigated with multiangle sensors (i.g. CHRIS Proba-1) or matching-resolution digital elevation models. Thus, automated classifiers should be applied with caution over large areas, ensuring robustness against these and other confounding factors.

### Caveats and limitations

The main result from model inversion at three sites was backed up by independent measurements in the field and previous studies (Sánchez-Azofeifa and Castro-Esau 2007, Mello et al. 2020). Separate measurements on leaf reflectance and in situ measurements of leaf traits and angles confirmed that lianas tend to have relatively flatter leaf angles and more cheaply constructed leaves with less dry matter and pigments. However, the model’s predictions were not always perfectly in line with ground truth data. On an absolute scale, ground measurements of mean leaf angles were lower than those predicted by PROSAIL2 (Appendix S6). One explanation for this is that we do not consider the effect of leaf clustering on stems and branches in our field campaign, while the PROSAIL2 result is more likely to reflect the integrated angles of leaves clustered on stems and branches. Direct measurements (i.e. not derived from remotely sensed data) on leaf angles, stems and branches collected from multiple sites would confirm the robustness of this result.

## Conclusions

Our work reveals a unique spectral signature for lianas at the canopy scale, detectable by aerial and spaceborne sensors. Models attribute the signature to lianas’ larger projected leaf area, lower leaf light absorption and higher transmission, leading to increased canopy-level light scattering. We observe the signal, characterized by higher forest albedo, in four distinct tropical sites, suggesting a consistent global pattern. If further confirmed, this could significantly impact our understanding of liana proliferation’s effects on forest energy budgets and understory dynamics. The identification of a general spectral signal for liana infestation across sites underscores the importance of developing accurate classifiers for quantifying liana abundance. Our successful use of radiative transfer models highlights the potential for creating robust, potentially physics-informed classifiers. In conclusion, fertile ground exists for future research into the liana spectral signal, its robust detection from remote sensing platforms, and the ecological consequences of liana proliferation.

## Acknowledgements

We thank Princeton University’s Carbon Mitigation Initiative, the European Research Council Grant 637643 (TREECLIMBERS), FWO (grants 1214720N/1214723N), and various Natural Environment Research Council grants [NE/P004806/1; NE/I528477/1; NE/L002604/1], along with the University of Nottingham Anne McLaren Research Fellowship. We acknowledge the Carnegie Airborne Observatory for their data contribution. Partial support came from the U.S. Department of Agriculture, Forest Service. The findings and conclusions in this study are those of the authors and do not necessarily reflect official positions of the USDA or U.S. Government.

