## Supporting information for "Why can we detect lianas from space?"

### Appendices: Why can we dectect lianas from space?

December 13, 2023

### S1 Datasets

Dataset numbering corresponds to table S1 and in the main text. References are supplied with a doi if they are not referenced in the main text. Full references are given if no doi exists.

Spectral datasets, location and sensor information

| | Scale | spatial extent* | Locations ** | Year | Species (vines) | Samples | platform | Sensor |
|---|---|---|---|---|---|---|---|---|
| 1 | Leaf | $< 10\ cm^2$ | Ispra, Italy | 1993 | 46 (3) | 330 leaves | Spectroradiometer | PerkinElmer LAMBDA-19 |
| 2 | | $< 10\ cm^2$ | Angers, France | 2004 | 43 (5) | 276 leaves | Spectroradiometer | ASD FieldSpec-FR |
| 3 | | $< 10\ cm^2$ | SLZ & PNM, Panama | 2004 | 40 (27) | 864 leaves | Spectroradiometer | Unispect PPSystems |
| 4 | | $< 10\ cm^2$ | SLZ & PNM, Panama | 2016 | 6 | 248 leaves | Spectroradiometer | SVC HR1024i |
| 5 | | $< 10\ cm^2$ | SLZ & PNM, Panama | 2017 | 28 | 852 leaves | Spectroradiometer | SVC HR1024i |
| 6 | | $< 10\ cm^2$ | CAM, CAR, FLMM, SB, UPR, Puerto Rico | 2017 | 11 | 104 leaves | Spectroradiometer | SVC HR1024i |
| 7 | | $< 10\ cm^2$ | TNF, Brazil | 2012 | 16 | 166 leaves | Spectroradiometer | ASD FieldSpec Pro |
| 8 | | $< 10\ cm^2$ | TNF, Brazil | 2017 | 11 | 759 leaves | Spectroradiometer | ASD FieldSpec Pro |
| 9 | | $< 10\ cm^2$ | VAR, C. & S. America | 2017 | 214(20) | 650 leaves | Spectroradiometer | ASD FieldSpec 3 & SVC HR1024i |
| 10 | | $< 10\ cm^2$ | DVCA, Malaysia | 2015 | 282 | 683 branches | Spectroradiometer | ASD FieldSpec 4 |
| 11 | | $< 10\ cm^2$ | KSFR, Malaysia | 2016 | 67 (20) | 492 leaves | Spectroradiometer | ASD FieldSpec Pro |
| 12 | Canopy | $1\ m^2$ | DVCA, Malaysia | 2014 | NA | 455 crowns | Airborne | Asia-Fenix |
| 13 | | $2\ m^2$ | BCNM, Panama | 2012 | NA | 544 crowns | Airborne | AToMS |
| 14 | Stand | $30\ m^2$ | NER,French Guina | 2006 | NA | 450 pixels | Satellite | Landsat |
| 15 | | $30\ m^2$ | NKM, Bolivia | 2001 | NA | 8164 pixels | Satellite | Hyperion |

Table S1.1: *Spectral sampling area is typically bewteen .6 to 3.5 $cm^2$ depending on leaf clip or artificial light specifications. **Abreviations alphabetically: Barro Colorado Nature Monument (BCNM), Cambalache State Forest (CAM), Carite State Forest (CAR), Danum Valley Conservation Area (DVCA), Fundacion Luis Munoz Marin (FLMM), International Institute of Tropical Forestry (UPR), Kabili-Sepilok Forest Reserve (KSFR), Nouragues Ecological Research Station (NER), Noel Kempff Mercado National Park (NKM), Parque Nacional San Lorenzo (SLZ), Parque Natural Metropolitan (PNM), Sabana (SB), Tapajos National Forest (TNF), and various sites throughout Central and South America (VAR).

#### S1.1 DATA SOURCES

1. (LOPEX), ECOSIS library: https://ecosis.org/package/leaf-optical-properties-experiment-database--lopex93-
2. (ANGERS), ECOSIS library: https://ecosis.org/package/angers-leaf-optical-properties-database--2003-
3. (Sánchez-Azofeifa et al 2009a): uploaded to dataserver upon acceptance.
4. (NGEE-TROPICS), LBL library: http://dx.doi.org/10.15486/ngt/1475180
5. (NGEE-TROPICS), LBL library: http://dx.doi.org/10.15486/ngt/1478523
6. (NGEE-TROPICS), LBL library: http://dx.doi.org/10.15486/ngt/1475180
7. (Wu et al. 2016), ECOSIS library: https://ecosis.org/package/2012-leaf-reflectance-spectra-of-tropical-trees-in-tapajos-national-forest
8. (Wu et al. 2017): uploaded to dataserver upon acceptance.
9. (Meireles et al. 2020): http://dx.doi.org/10.6084/m9.figshare.12449147.v1
10. (Nunes et al 2019, doi: 10.1088/1748-9326/ab2eae): uploaded to dataserver upon acceptance.
11. (Sepilok Forest Reserve spectral data): uploaded to dataserver upon acceptance.
12. (Chandler et al 2020), NERC library: http://dx.doi.org/10.5285/c708cad9950c45b1af0d4e9ca944f09a
13. (Marvin et al 2016), uploaded to dataserver upon acceptance.
14. (Tymen et al 2016): data available from DRYAD: https://doi.org/10.5061/dryad.1pc19
15. (Foster et al 2006), EO-1 Hyperion data: https://doi.org/10.5066/P9JXHMO2
16. (Rodríguez-Ronderos, et al 2016): uploaded to dataserver upon acceptance.
17. (Visser and Detto), leaf angle data: uploaded to dataserver upon acceptance.

#### S1.2 DATA DESCRIPTIONS

- **Dataset 1 - 2**: Both datasets 1 and 2 are well known in the remote sensing community as the LOPEX and ANGERS datasets respectively. Leaf biochemical and spectral measurements were conducted for both datasets.

  For LOPEX, a total of 330 leaves from 46 species were collected in 1993 at the Joint Research Centre at Ispra in Italy. The optical measurements were performed on the leaf blades using a Perkin Elmer Lambda 19 double-beam spectrophotometer. The ANGERS dataset comprised 276 leaves from 43 species collected in June 2003 at the INRA Agricultural Experiment Station of Angers in France. In this case, the optical measurements were conducted using a FieldSpec-FR portable Spectroradiometer. In both datasets, an integrating sphere coated with BaSO4 was used, enabling the measurement of reflectance and transmittance spectra within the range of 350 to 2500 nm.

  Following the spectral measurements, leaf discs with a diameter of 5 mm were sampled to determine biochemical content. Pigments were extracted by grinding the leaf discs in a chilled mortar and using 95% ethanol (v/v). Water content and dry matter content ($g.cm^{-2}$) were calculated by weighing and averaging three discs with a diameter of 10 mm from fresh and dry samples. The fresh weight was measured before placing the leaf samples in a drying oven at 85 $\circ$ C for 48 hours. After drying, the samples were reweighed to determine the dry weight and water content.

  Additional details on the LOPEX dataset can be found in (Hosgood et al., 1994)[1], and for the ANGERS dataset, refer to Pavan et al. (2004)[2].

- **Dataset 3**: During the rainy season in August 2004, sun leaves from ten individuals of the most abundant species within the crane arm's reach were collected. The sampling followed established protocols, resulting in a total of 26 liana and 9 tree species sampled from PNM, and 9 liana and 9 tree species from FS. Each species had a maximum of 10 leaves collected, in accordance with Panamanian National Environment Authority guidelines. The leaves, free of epiphytes, mosses, and galls, were immediately stored in sealable plastic bags with moist paper towels and then placed in a larger ice-containing black plastic bag.

  Spectral reflectance measurements (400 to 1100 nm) were taken on the day of collection using a portable spectrometer. This method captures leaf structure, pigment sizes, and water content but not rapid photochemical changes. A bifurcated fiber optic and leaf clip were used, and the data included visible/near-infrared reflectance. Leaf cores were then preserved

[1] Hosgood, B., Jacquemoud, S., Andreoli, G., Verdebout, J., Pedrini, G., Schmuck, G., 1994. Leaf Optical Properties EXperiment 93 (LOPEX93). Eur. Comm. Jt. Res. Centre, Inst. Remote Sens. Appl. Rep. EUR 16095 EN 11.

[2] Pavan, G., Jacquemoud, S., De Rosny, G., Rambaut, J.-P., Frangi, J.-P., Bidel, L.P.R., 2004. RAMIS: A NEW PORTABLE FIELD RADIOMETER TO ESTIMATE LEAF BIOCHEMICAL CONTENT.

for pigment analysis. Additionally, diffuse transmittance and reflectance were measured for five leaves per species, with absorptance estimated as the complement of these values. These data were aggregated for further analysis at the structural group level.

Leaf thickness was measured on five mature leaves from different individuals of each species. The average thickness was calculated from six points near the apex, middle, and base using a leaf thickness micrometer. These measurements were made within two hours of collection. Additionally, the fresh and dry weights of the leaves were determined, with dry weight measured after drying at 60 °C for 36 hours. Nitrogen and phosphorus concentrations were estimated using a combustion elemental analyzer and colorimetric analysis, respectively. Specific leaf area (SLA) was calculated as the ratio of fresh surface area to dry weight.

Detials can be found in Sánchez-Azofeifa *et al.* 2009.

- **Dataset 4, 5 & 6**: A total of 27 canopy tree species from two sites in Panama were selected for intensive field measurements of leaf reflectance spectra and traits in 2016 and 2017.

  The sampled leaves were categorized into two main age classes: immature leaves (less than 2 months old), which encompassed leaves from emergence to fully expanded but not fully green, thickened, or physiologically matured, and mature leaves (2 months or older). This age classification aligns with similar categories proposed by Coley (1983, doi:10.2307/1942495) and Wu et al. (2016,doi: 10.1111/nph.14051). However, unlike Wu et al. (2016), where three age categories (young, mature, and old) were used, our study combined the mature and old age classes into a single mature age class.

  Field measurements were conducted during the dry seasons of 2016 and 2017 on sunlit upper canopy foliage in Panama. In the 2016 field campaigns, which took place in mid-February and mid-April, we sampled the dominant leaf class(es) from eight trees at each site. In February 2017, the measurements included both age classes if they were present within the top meter of a sunlit branch from four canopy tree species at the SLZ site.

  Leaf reflectance at the Panamanian sites was measured using a Spectra Vista Corporation (SVC) HR-1024i spectroradiometer (SVC, Poughkeepsie, NY, USA). The spectral range of the instrument was 350-2500 nm, with a spectral resolution of 3.5 nm at 700 nm, 9.5 nm at 1500 nm, and 6.5 nm at 2100 nm. The measurements were conducted with the SVC LC-RP-Pro foreoptic. Details given in Serbin *et al.* (2019, doi: 10.1111/nph.16123)

- **Dataset 7 & 8**: Measurements of leaf full-spectrum reflectance and transmittance were conducted across 54 tropical tree species, covering the wavelength range of 350-2500 nm. The dataset includes leaves collected from both fully sunlit and shaded canopy strata, as well as leaves representing different age categories, including young, mature, old, and senescent leaves. For each sample, information on the relative age estimate, leaf canopy position, and sample number is provided.

The data was collected as part of the 2017 NGEE-Tropics / NASA G-LiHT airborne campaign. Additional datasets related to this study contain details such as sample photographs, leaf traits (e.g., leaf mass per area (LMA), water content), and leaf carbon and nitrogen measurements.

Leaf reflectance was measured using a Spectral Evolution PSR+ full-range spectroradiometer. To perform the measurements, a custom fiber optic connected to a Spectra Vista Corporation LC-RP-Pro foreoptic was utilized. More detauls given in Wu *et al.* (2016,doi: 10.1111/nph.14051) and Wu *et al.* (2017).

- **Dataset 9**: Meireles et al. (2020) compiled a comprehensive dataset comprising over 16,000 leaf-level reflectance spectra ranging from 400 to 2400 nm. This dataset encompassed 544 seed plant species found in temperate and tropical regions across the Americas and Europe (Fig. 1 in maintext). The included spectra were solely derived from mature, sun-exposed leaves measured during the spring or summer seasons. In our study, we utilized the aforementioned dataset, focusing on a subset of 650 leaves from 214 tropical trees and lianas identified in the dataset using tables provided by Martin and Asner (2008).

  The data contains leaf spectra, obtained from two full-range field spectroradiometers: an ASD FieldSpec 3 (Analytical Spectral Devices, Boulder, CO, USA) and an SVC HR-1024i (Spectra Vista Corp., Poughkeepsie, NY, USA) with measurements were conducted using leaf clips and artificial light sources. Processing details are given in Meireles et al. (2020), and include harmonization among sensors.

- **Dataset 10**: In all field surveys a random selection of five leaves attached to the branches was made, excluding damaged and young leaves. Reflectance spectra spanning the range of 350-2500 nm were acquired using a FieldSpec 4 spectroradiometer, manufactured by Analytical Spectral Devices (ASD) based in Boulder, Colorado, USA.

  To ensure accurate measurements, the spectroradiometer's contact probe was securely mounted on a clamp and pressed firmly against the sample, while a black background was employed to eliminate any external light interference. The spectral measurements were taken at the midpoint between the main vein and the leaf edge, approximately halfway between the petiole and the leaf tip. The abaxial surface of the leaf faced towards the probe during measurement.

  To maintain calibration, readings were periodically adjusted against a Spectralon white reference panel after every five samples. In all subsequent statistical analyses, the spectral data were trimmed to the 400-2500 nm range, and the mean reflectance values from the five spectra per branch were utilized. More details in Nunes *et al.*(2019, doi: `10.1088/1748-9326/ab2eae`)

- **Dataset 11**: Nine 4-ha plots were established in Sepilok between 2000 and 2001. The most recent survey in 2014 included the measurement and identification of all trees with a diameter at breast height (DBH) of 5 cm or more, with 91% identified to the species level. In a 0.25-hectare subplot within each plot, liana abundance, biomass, and diversity were measured for stems over 0.5 cm DBH. Sampling of tree and liana branches occurred in April-May 2016, resulting in 54 tree and 23 liana species sampled from 179 individuals. Common tree species accounted for over 50% of the total basal area, while the sampled liana species represented over 50% of the liana basal area.

  To sample sunlit branches, researchers collected small branches from accessible trees and lianas in the forest canopy, including neighboring crowns of less abundant species for larger trees. The branches were immediately cut underwater, preserved in a water-filled plastic enclosure, and stored in a closed Styrofoam box with ice packs. Bulk leaf samples were collected and dried for elemental analysis. Measurements of traits were conducted in close proximity to the forest edge at the Phytochemistry Laboratory of the Sabah Forest Research Centre.

  For each collected branch, three clean leaves were selected for hyperspectral reflectance measurements. The branch was hydrated overnight, and the weight of three leaves was measured the next morning. The leaves were scanned for area determination and then dried in an electric oven. Elemental concentrations and stable isotope ratios were analyzed using a Costtech Elemental Analyser attached to a Thermo DELTA V mass spectroradiometer.

  Hyperspectral reflectance signatures were measured using a FieldSpec Pro spectroradiometer equipped with a contact reflectance probe and an integrated light source. The spectroradiometer employed three sensors with shared optics, covering different wavelength ranges. To ensure accurate measurements, leaf samples were placed on a low reflectance black background, and a specific procedure was followed to eliminate stray light and account for drift in the illumination source and sensor using dark current and white reference measurements (Chavana, 2017, doi: 10.1111/nph.13853).

  **Dataset 12**: In November 2014, the UK Natural Environmental Research Council's Airborne Research Facility (NERC-ARF) collected manned airborne hyperspectral data over a primary forest area of approximately 2083 ha. The data were acquired using a Dornier 228-201 aircraft, flying at speeds ranging from 127 to 139 knots at altitudes of 2335-2429 m. A total of 10 flightlines were flown, capturing detailed information about the forest (Chandler, 2020).

Hyperspectral imagery was collected using a FENIX sensor with a spatial resolution of 3 m. The data were calibrated in collaboration with the NERC Field Spectroscopy Facility. Radiometric corrections were applied to the dataset, removing bands with missing or overly saturated data. ENVI FLAASH Atmospheric Correction was used for atmospheric correction. To address variations in reflectance values between flightlines, spectral values were adjusted based on the difference in reflectance between overlapping pixels from adjacent flightlines. A Standardized Principal Component Analysis (SPCA) was performed to reduce the dimensionality of the data, retaining eight principal components that explained over 99% of the variation.

Two datasets were collected for the liana canopy cover survey: ground-based below-the-canopy data and UAV-based above-the-canopy data. Ground data were collected between 2017 and 2019, involving the visual delineation of individual tree crowns using LiDAR data uploaded to a tablet computer connected to a GPS receiver. The percentage of liana infestation within each tree crown was estimated independently by field teams through mutual agreement. UAV-derived imagery was used to collect liana infestation data at the tree-level and hyperspectral pixel-scale. A DJI Phantom 4 Advanced quadcopter with an integrated RGB camera was employed for data acquisition, with geo-tagged images processed to generate a single orthorectified image. Individual tree crowns were identified, and liana infestation was estimated by visually assessing the percentage of a pixel covered by lianas within crown boundaries. For additional information, please refer to Chandler (2020).

**Dataset 13**:Airborne collection of imagery over the Gigante Peninsula in Barro Colorado National Monument in the Republic of Panama occurred in the dry season of February 2012. The Carnegie Airborne Observatory Airborne Taxonomic Mapping System (AToMS) acquired high-resolution data of the site with an integrated full-spectral range (visible-to-shortwave infrared) imaging spectroradiometer (Asner et al., 2012, doi: 10.1016/j.rse.2012.06.012). The AToMS visible-to-shortwave infrared imaging spectroradiometer (VSWIR) measures spectral radiance in 481 contiguous channels spanning the 252-2648 nm wavelength range. At a flight altitude of 2000 m, the VSWIR data collection provided a ground sampling distance of 2.0 m throughout the study landscape. VSWIR spectroradiometer data were radiometrically corrected from raw digital number (DN) values to radiance (W $sr^{-1}$ $m^{-2}$ $nm^{-1}$) using a flat-field correction, radiometric calibration coefficients, and spectral calibration data collected in the laboratory, with further post-processing described in detail by Asner et al. (2012). Reflectance imagery was corrected for cross-track brightness gradients using a bidirectional reflectance distribution function. Individual tree crown field assessments were collected in July and August of 2013. Further details on imagery and field data acquisition and processing can be found in Marvin et al. (2016).

- **Dataset 14**: Tymen et al (2016) supply the georeferenced locations of forests heavily infested by lianas (hereafter “liana forest”) which was delineated using lidar, aerial photography and a ground census. We obtained a georeferenced 30-m resolution Landsat Thematic Mapper image (L1TP) acquired on October 8 2006 of the study area. The image was atmospherically corrected using the FLAASH algorithm in ENVI 5.5.

- **Dataset 15**: The obtained Hyperion image underwent several checks and analyses. Firstly, a minimum noise fraction transform (MNF) was performed to detect smile, resulting in 25 MNF bands based on eigenvalues greater than or equal to 3 (Goodenough et al., 2003, doi: `10.1109/TGRS.2003.813214`). Smile was observed in the first two MNF bands, noticeable through the brightness gradient (Goodenough et al., 2003). To mitigate the inherent ”striping” in the Hyperion data and address the smile effect, a ”destreaking” algorithm was applied. This algorithm normalized the pixel values in each sample to the overall mean and standard deviation of all samples (Datt et al., 2003, doi: `10.1109/TGRS.2003.813206`). Before destreaking, specific areas such as clouds, cloud shadows, and background pixel values resulting from VNIR and SWIR coalignment were masked in the 196-band image. Masks were generated using pixel value thresholds in three bands to exclude values that could bias the destreaking results. The MNF transform was then applied to the destreaked image, resulting in a reduction in striping and brightness gradient compared to the ”undestreaked” image.

  Subsequently, atmospheric correction was performed using ACORN (version 5, mode 1.5pb) on the destreaked 196-band image. ACORN is designed specifically for pushbroom sensors like Hyperion, which exhibit cross-track spectral calibration variation causing smile. ACORN utilizes a wavelength and full-width half-maximum (FWHM) array file generated from the center wavelength and bandwidth datasets in the HDF (L1R) file. The output of the atmospheric correction process was an atmospherically corrected image, transforming the radiance values to apparent surface reflectance.

- **Dataset 16**. The contribution of lianas to Plant Area Index (PAI) was estimated in a liana removal experiment in the secondary forest of Gigante Peninsula, Barro Colorado Nature Monument (Republic of Panama). In 2008, sixteen 80m x 80m plots were established and all trees and lianas $\geq$ 1cm DBH were measured in the central area of each plot (60m x 60m). In 2011, immediately after a census of lianas and trees, all lianas were cut in eight randomly selected forest plots, with the remaining eight plots serving as non-manipulated control plots.

  Mean per-plot Plan Area Index (PAI) were estimated as the sum of Leaf Area Index (LAI) and Wood Area Index (WAI) with the the Li-Cor LAI-2000 plant canopy analyzer (Li-Cor Biosciences, Lincoln, NE, USA). PAI measurements were taken along a grid of 49 points within the central 60 m x 60m of each plot, 50 cm and 1 m above the soil surface (n=98 measurements per plot). Simultaneously, open-sky measurements of PAI were taken every 30 seconds with a second Li-Cor LAI-2000 outside of the forest canopy on the edge of Lake Gatun. PAI measurements were restricted to the northern half of the sensor (180 degree cap) for both open-sky and within-forest measurements to ensure that the open-sky measurements did not intercept forest leaf area, and the within-forest measurements did not include the shadow of the operator. Measurements of PAI were taken fifteen days before the liana cut in 2011 in all sixteen plots, one month after the liana cut, and every consecutive year for four years. The vast majority of lianas had fallen one year after the liana cut. At every sampling period, plots were measured in the same order, ensuring consistency and comparable measurements. Plant area index (PAI) measurements were processed with the Li-Cor FV2000 Analysis Software (2005, Li-COR, Biosciences, Lincoln, NE, USA). Further details are in Rodríguez-Ronderos et al. (2016).

*Calculation of $f_2$ using Li-Cor data*. A naive but straightforward method to estimate $f_2$ would involve dividing the average PAI of the removal plots after liana cutting by the average PAI of the control plots, represented as $(P_{rem}/P_{con})$. However, this approach can introduce bias because not every tree is affected by lianas, and the extent of infestation varies among trees. Particularly, when fewer trees in a plot are infested by lianas, this estimator tends to be more biased. To address this, we derive an unbiased estimator of $f_2$ below.

Past research indicates that liana leaves displace tree leaves in a 1:1 ratio (Kira & Ogawa, 1971[3]). This observation is consistent with dataset 15, which revealed that the PAI post-liana removal matched the PAI in control plots four years post-cutting, as discussed in depth in Rodriguez-Ronderos et al. (2016). This consistency implies that the maximum achievable plot PAI, without lianas, closely resembles that in the control plots. Given this empirical finding, we posit that the expected maximum PAI without lianas ($P_{max}$) equals the observed PAI in the control plots ($P_{con}$).

The reduction in total PAI due to lianas is then tied to: 1) the fraction of infested trees in the plot ($I$), 2) the average tree infestation level ($L$), and 3) the portion of liana leaves in a fully infested canopy, given as 1-$f_2$ when L=1. Thus, the expected PAI in a control plot is the sum of both liana and tree leaves:

$$P_{con} = I[L(1 - f_2)P_{max} + [1 - L(1 - f_2)]P_{max}] + (1 - I)[P_{max}] = P_{max} \quad (1)$$

[3]Kira T, Ogawa H (1971) Assessment of primary production in tropical and equatorial forests. In: Duvigneaud P (ed) Productivity of forest ecosystems. Proc Brussels Symp Unesco, Paris

This equation breaks down the total PAI in a plot into contributions from infested (I) and uninfested trees (1-I). When a tree is infested, the maximum PAI is subdivided into liana and tree components using $f_2$ (the fraction of tree leaves) and the infestation level (L). For instance, the portion of liana leaves in an infested crown is given by $L(1-f_2)$. As predicted, Eq. 1 simplifies to $P_{con} = P_{max}$, mirroring empirical findings. Conversely, the PAI after liana removal is expressed by:

$$P_{rem} = I[P_{max}[1 - L(1 - f_2)]] + (1 - I)[P_{max}] = P_{max}[I(1 + f_2L - L) + (1 - I)] \quad (2)$$

This doesn't simplify to $P_{max}$: $P_{rem} < P_{max}$ whenever $L(1 - f_2) > 0$. (i.e., when lianas add to the PAI), consistent with empirical observations. Knowing I and L, we can then solve for $f_2$ as:

$$f_2 = \frac{P_{max}(IL - 1) + P_{rem}}{P_{max}IL} \quad (3)$$

Given average values of I and L for the plots (0.84 and 0.65, respectively, as per van der Heijden et al. 2016 and S. Schnitzer, personal communication), $f_2$ can be deduced using the observed $P_{rem}$ - and presuming $P_{con} = P_{max}$.

- **Dataset 17**. Leaf angle data, which we obtained by collecting and analyzing leveled digital photographs, is detailed in the main text.

### S2 Hierarchical models, MCMC sampling and prior distributions

We fit leaf-level RTMs to each individual leaf spectra using the described MCMC method in the maintext and obtained RTM model parameters for each leaf (N, $C_{ab}$, $C_{ar}$, $C_{br}$, $C_{an}$, $C_w$, $C_m$). Subsequently, we employed Bayesian hierarchical models to to analyze variance across growth forms, adhering to the data's hierarchical structure: leaf measurements within species, and species within studies. Generally, the leaf-level model took the following form:

$$T_{ijk} = \beta_0 + \beta_i + \beta_{jk} + \beta_k + \epsilon \quad (4)$$

where $T_{ijk}$ represents one of the RTM model parameters mention above, fit for growth form i (liana or tree) from species j nested in study k. The beta parameters had normal priors with uninformative hyperpriors (following Gelman 2004[4]):

$$\beta_n \sim N(0, \sigma_n) \quad (5)$$

[4]Gelman, 2004, Prior distributions for variance parameters in hierarchical models,EERI Research Paper Series No 6/2004

where n represents one of hierachical grouping factors such as species, site or growth form.

For canopy and stand scales, we fitted the PROSAIL2 model for each site individually, without using hierarchical models to break down variance among growth forms. This is because PROSAIL2 was directly adapted to simulate liana-infested crowns/pixels as two-layer canopies with separate vertical sections (defined by $f_2$ and $L$), each characterized by growth form-specific optical parameters (see maintext). Hence, a Bayesian fit of our PROSAIL2 model already break down variance among growth forms.

We used predicted growth-form level means and variances across species and sites as prior information for leaf parameters (since the same leaf RTM is paired with the canopy-RTM). The prior possesses the mean trait value for growth-form $i$, allowing for variance from species, study, and other unexplained sources. This led to a largely overlapping prior distributions, with specific absolute values detailed in table S2.2.

We employed Markov-chain Monte Carlo (MCMC) methods to fit the model. Specifically, we established the posterior distribution for a vector of RTM parameters, denoted as a vector $\boldsymbol{\phi}$, and a vector $\mathbf{R}$ of observed reflectance values.

$$P(\boldsymbol{\phi}, \sigma \mid \mathbf{R}) \propto P(\mathbf{R} \mid \boldsymbol{\phi}, \sigma) P(\boldsymbol{\phi}) P(\sigma) \tag{6}$$

Here P($\mathbf{R} \mid \boldsymbol{\phi}, \sigma$) is the likelihood of observing reflectance vector $\mathbf{R}$, which is assumed to be normally distributed around the simulated reflectance predicted from parameter set $\boldsymbol{\phi}$ with a variance of $\sigma^2$. P($\boldsymbol{\phi}$) and P($\sigma$) are the prior probabilities of parameters $\boldsymbol{\phi}$ and $\sigma$.

Each model inversion was initialized each using randomly drawn parameter values from the prior distributions (see table S2.1 below), and we ran the algorithm for ten independent chains until convergence. We used a burn-in of 10,000 and 100,000 iterations for leaf scale and larger scales, respectively. MCMC convergence was determined based on a value of the Gelman-Rubin multivariate potential scale reduction factor of less than 1.035 as earlier explorative analyses revealed that this cutoff improved the fit beyond the standard 1.05 cutoff (Gelman & Rubin, 1992,doi: 10.1214/ss/1177011136). Convergence typically occurred after 10,000 to 10,000,000 iterations, after burn-in, depending on model complexity. Each chain was run in parallel on a high-performance computing cluster, and we implemented a thinning interval of 10, storing only every tenth iteration, as this ensured both 1) an accurate and representative sample of the joint posterior distribution and 2) saved memory usage tenfold. After applying the burn-in and thinning filter, credible intervals (CI) were calculated as the 2.5 and 97.5 percentile of the marginal distributions of each parameter - which are estimated by sampling from the joint posterior distribution using the MCMC.

Model parameters priors in leaf level analysis (PROSPECT)

| Trait | parameter | Prior | a | b |
|---|---|---|---|---|
| Leaf stuctural parameter | N | Uniform | 1 | 5 |
| Chlorophyll A & B content per unit leaf area | $C_{ab}$ | Uniform | 0 | 200 |
| Carotenoid content per unit leaf area | $C_{ar}$ | Uniform | 0 | 50 |
| Water mass per unit leaf area. | $C_w$ | Uniform | 0 | 0.1 |
| Leaf dry matter per area | $C_m$ | Uniform | 0 | 0.1 |
| Anthocyanin content | $C_{an}$ | Uniform | 0 | 50 |
| brown pigments content | $C_{br}$ | Uniform | 0 | 10 |
| Error | $\sigma$ | Uniform | $1 \times 10^{-9}$ | 0.1 |

Table S2.1: Prior distributions used to fit PROSPECT5,5b and D to leaf reflectance data. Columns a & b refer to the upper and lower bounds of the uniform distribution.

Stand and canopy scale model priors (PROSAIL2) for the top (1) and bottom (2) layer.

| Trait | parameter | Prior | a / $\mu$ | b / $\sigma$ |
|---|---|---|---|---|
| Leaf stuctural parameter | $N_1$ | Normal | 1.72 | 0.27 |
| Chlorophyll A & B / area | $C_{ab,1}$ | Normal | 55.65 | 33.89 |
| Carotenoid content / area | $C_{ar,1}$ | Normal | 10.18 | 4.06 |
| Water mass / area | $C_{w,1}$ | Normal | 0.011 | 0.0056 |
| Leaf dry matter / area | $C_{m,1}$ | Normal | 0.0068 | 0.0021 |
| Anthocyanin content / area | $C_{an,1}$ | Uniform | 0 | 20 |
| brown pigments content | $C_{br,1}$ | Normal | 0.20 | 0.23 |
| Leaf stuctural parameter | $N_2$ | Normal | 1.86 | 0.58 |
| Chlorophyll A & B / area | $C_{ab,2}$ | Normal | 65.54 | 32.77 |
| Carotenoid content / area | $C_{ar,2}$ | Normal | 14.56 | 7.14 |
| Water mass / area | $C_{w,2}$ | Normal | 0.019 | 0.012 |
| Leaf dry matter / area | $C_{m,2}$ | Normal | 0.01 | 0.0059 |
| Anthocyanin content | $C_{an,2}$ | Uniform | 0 | 20 |
| brown pigments content | $C_{br,2}$ | Normal | 0.11 | 0.14 |
| Leaf Area Index | $LAI$ | Uniform | 0 | 15 |
| Leaf inclination angle | $\Omega$ | Uniform | 0 | 90 |
| Fraction secondary particles | $f_2$ | Uniform | 0 | 1 |
| Canopy dissociation factor | D | fixed | 1 | 1 |
| hotspot | $\kappa$ | Uniform | 2.94e-05 | 0.04 |
| Verticle crown cover fraction | Cv | fixed | 1 | NA |
| Crown shape | $\zeta$ | fixed | site specific | NA |
| Solar zenith angle | $\theta_s$ | fixed | site specific | NA |
| Observer zenith angle | $\theta_o$ | fixed | site specific | NA |
| Sun-sensor relative azimuth angle | $\psi$ | fixed | site specific | NA |
| Error | $\sigma$ | Uniform | $1 \times 10^{-9}$ | 0.15 |

Table S2.2: Prior distributions used to fit PROSAIL2 to leaf reflectance data. Columns a / $\mu$ & b / $\sigma$ refer to the upper and lower bounds of the uniform distribution or the mean and standard deviation of the normal distribution respectively. Whenever the value was fixed, the fixed value is reported in the a / $\mu$ column. Site specific parameters are taken from the scene metadata where appropriate (e.g. observer, sun and sensor angles). The prior range for the $\kappa$ parameter was estimated from the min and max values calculated across all sites using published global datasets on canopy height (Healey et al. 2015, `http://dx.doi.org/10.3334/ORNLDAAC/1271`) and leaf sizes (Wright et al. 2017 Science, 357).

### S3 Model robustness analyses

To get an idea of the robustness of various aspects of our inversion approach we conducted several simulation studies. Following Shiklomanov et al (2016), we identify three potential interacting error sources in our modeling study that could affect our results.

1. Model formulation error, which refers to a lack of fit, and overprediction or underprediction. We guard against model formulation error threefold. First, we evaluate the fit of each model by comparing how well the model can reproduce the observed spectral reflectance (as quantified by $R^2$). Second, for canopy and stand scales, we test PROSAIL2's ability to capture the difference between liana and tree signals in an unbiased fashion (e.g. on the 1:1 line). However, goodness of fit to data alone is not a sufficient test of a complex model as many free parameters can provide too much flexibility (e.g. White and Marshall 2019, doi: 10.1016/j.tree.2019.01.006). Therefore, finally, we compared whether inversely estimated differences between lianas and trees could be verified with independent measurements of the traits represented by the inverse parameters.

2. Parameter identifiability, or whether all parameters can be estimated from data. Lack of identifiability may indicate problems with the inversion procedure. We evaluated the role of parameter identifiability by simulating spectral signals from models parameterized with random parameters, and then back fit the same models using our Bayesian inversion algorithm. This analyses revealed that all free parameters were reasonably to well retrievable, i.e. with the generating parameter falling within 95% CI of the atleast 70% of the time given our prior settings.

3. Measurement error in spectra and traits (i.e. signal to noise ratios). Measurement error may systematically bias parameter estimates and lead to false differences between groups whenever groups differ in the degree of error (see e.g. Detto et al., 2019, doi: 10.1111/ele.13372). In our case, measurement error proved difficult to directly deal with because our datasets lack the repeated measurements needed to quantify measurement error. We therefore simulated the effect of increasing error in reflectance spectra to see if any differences in inversely estimated parameters could be explained by a differing error rate between groups. We found no consistent trends in how parameters responded to error in spectral data when the error rate differed among groups. Measurement error on verification traits, on the other hand, is expected to have a conservative effect on our results, i.e. it should lead to regression dilution, or a decreased ability of the model to predict physical traits from reflectance data.

### S4 Leaf reflectance data and leaf model fits

#### S4.1 Leaf reflectance data

The figure below (Fig. S4.1) shows observed leaf reflectance for five plant groups, and Kullback-Leiber Divergence (KLD) which is a measure of the (theoretical) ability of classifiers to discriminate between plant groups as measured by the observed spectral distributions over wavelengths. The panel in column A, shows the measured reflectance in the solar spectrum (wavelengths between 400 to 2500 nm) of various plant species, grouped in broad categories. Here, the black dashed line indicates the mean reflectance for lianas, while the colored envelopes represent the 2.5% to 97.5% percentile interval of measured reflectance for each group. The groups are, from top to bottom, crops species, herbs and grasses, shrubs, trees and lianas (see also color legend in column C). The panel in column B, shows the distribution of KLD scores at each wavelength for each group compared to the liana distribution (bottom Panel in A). Here, KLD scores were bootstrapped at equal sample size between groups and, for lianas, within the group. The panel in column C, shows the average total divergence across all wavelengths and includes the bootstrapped 95% confidence intervals (dots and whiskers). Together the panels show that leaf reflectance compared across plants shows large variation, and substantial overlap, cautioning that dectection algorithms could confuse groups in a global analysis. In the KLD calculation, we accounted for variance sensitivity to sample size. We ensured equal sample sizes by bootstrapping at the smallest common sample size for 10,000 iterations for each group. Groups with at least 50 spectral samples were included as the expected relative error for $\sigma(\lambda)$ is then $<10\%$. The bootstrap samples across inter- and intraspecific and site-specific variance for all plant groups. Hence, detectable significant differences imply robust distributional divergence – detectable to a classifcation algotithm without any additional information (i.e. phylogeny)

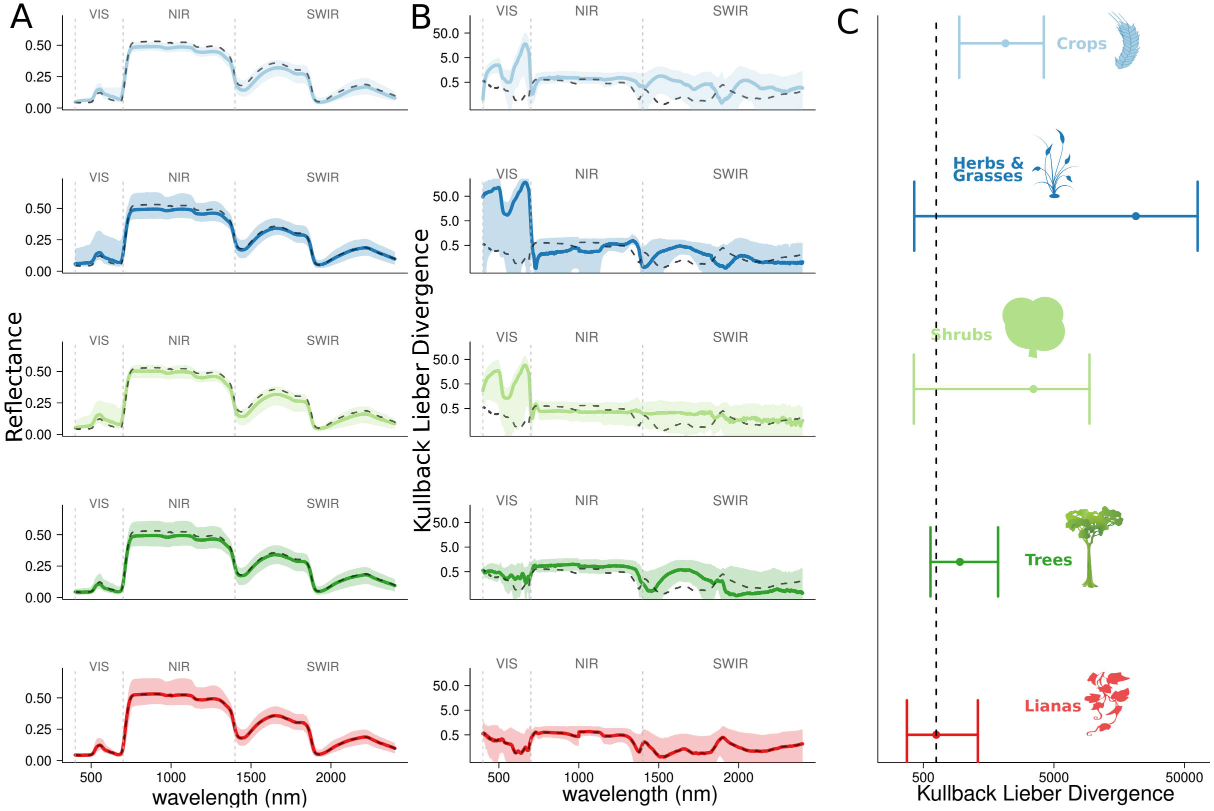


Figure S4.1: See above.

#### S4.2 Summary of leaf model fit statistics

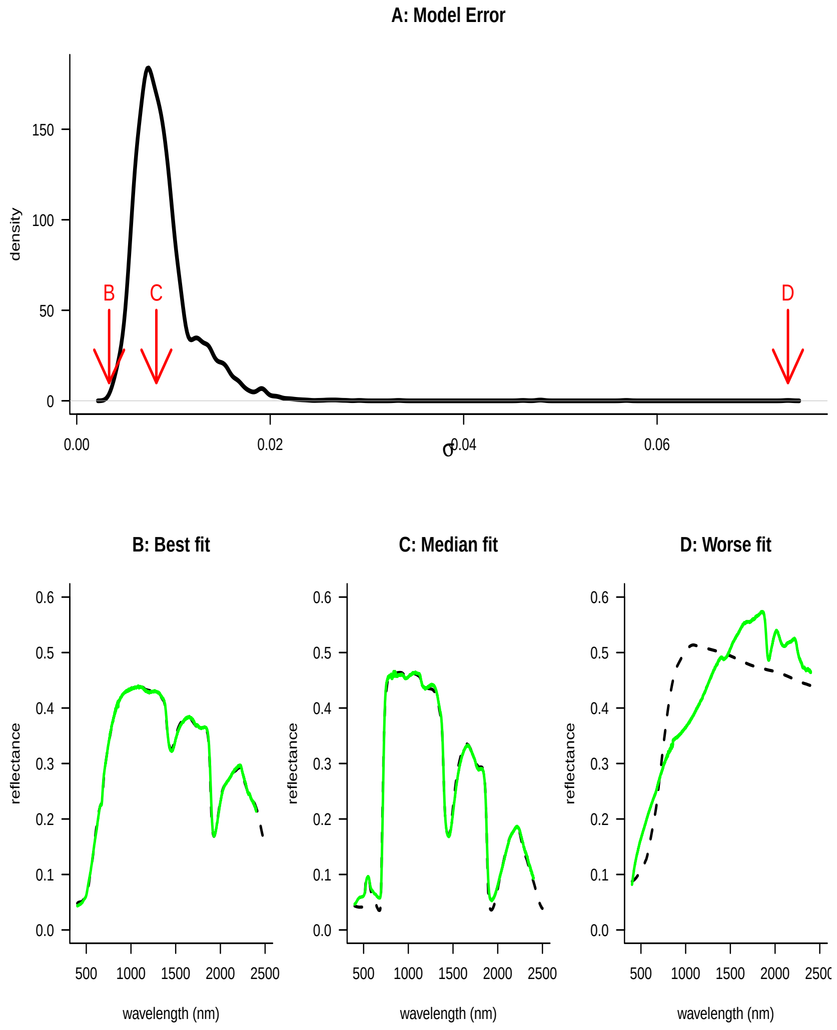


Figure S4.2: Leaf model fit statistics for the PROSPECTD inversion for 5424 leaves. Panel A gives the distribution of model errors ($\sigma$). In A red arrows indicate the position of the following three panels that represent the best fit (B), median fit (C) and worse fit (D). In B - D, black lines indicate the model fit, while green lines indicate the observed reflectance.

### S5 Posterior distributions

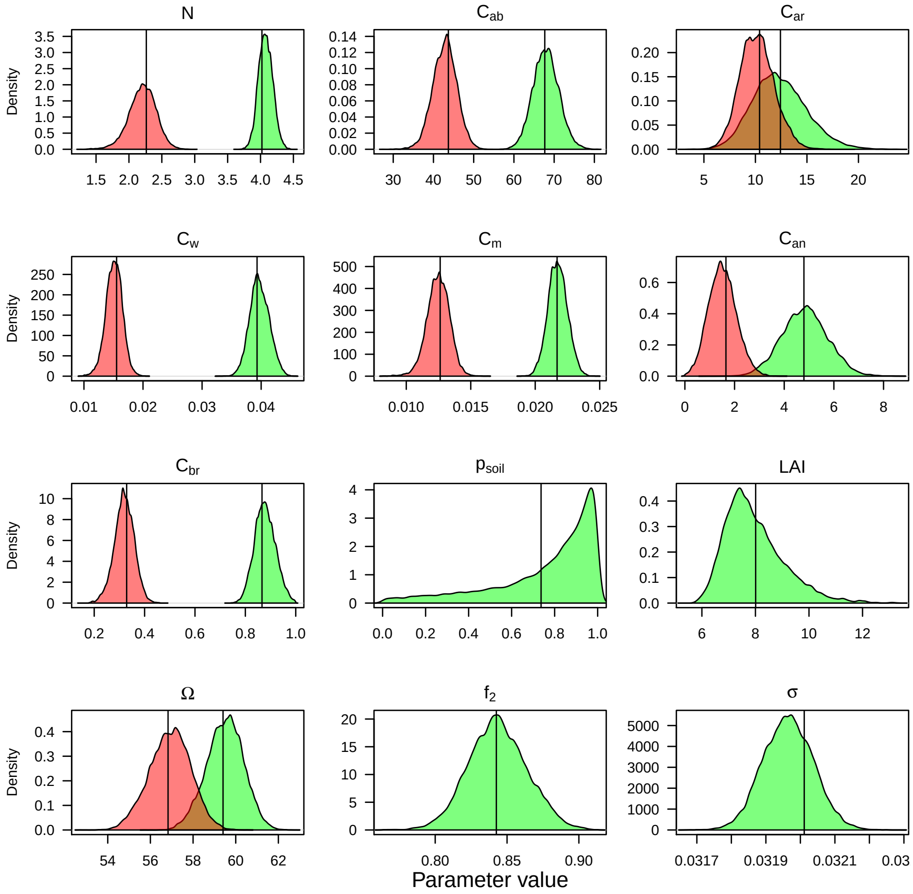


Figure S5.1: Marginal density plots for the posterior distributions for each model parameter of the PROSAIL2 model fit to the Bolivia dataset. Parameters that differ between lianas and trees are given in red and green respectively. Vertical lines indicate mean values for all distributions.

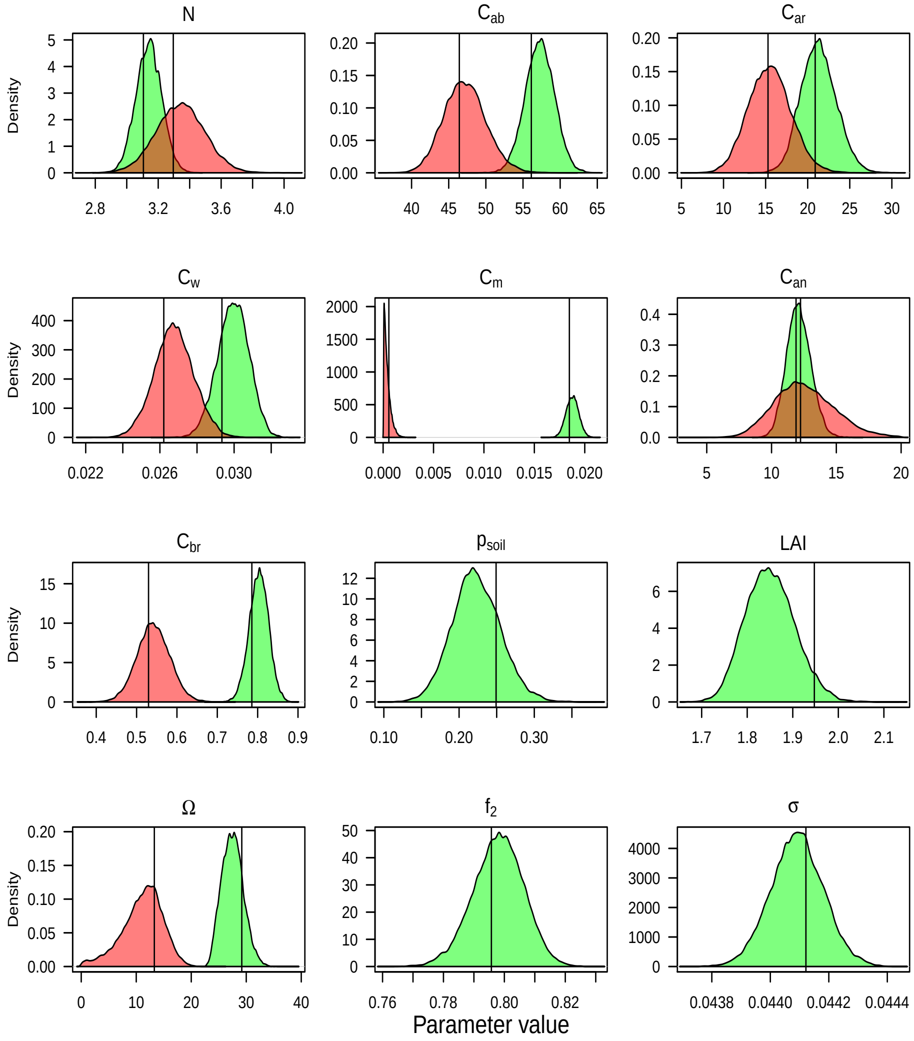


Figure S5.2: Marginal density plots for the posterior distributions for each model parameter of the PROSAIL2 model fit to the Malaysian dataset. Parameters that differ between lianas and trees are given in red and green respectively. Vertical lines indicate mean values for all distributions.

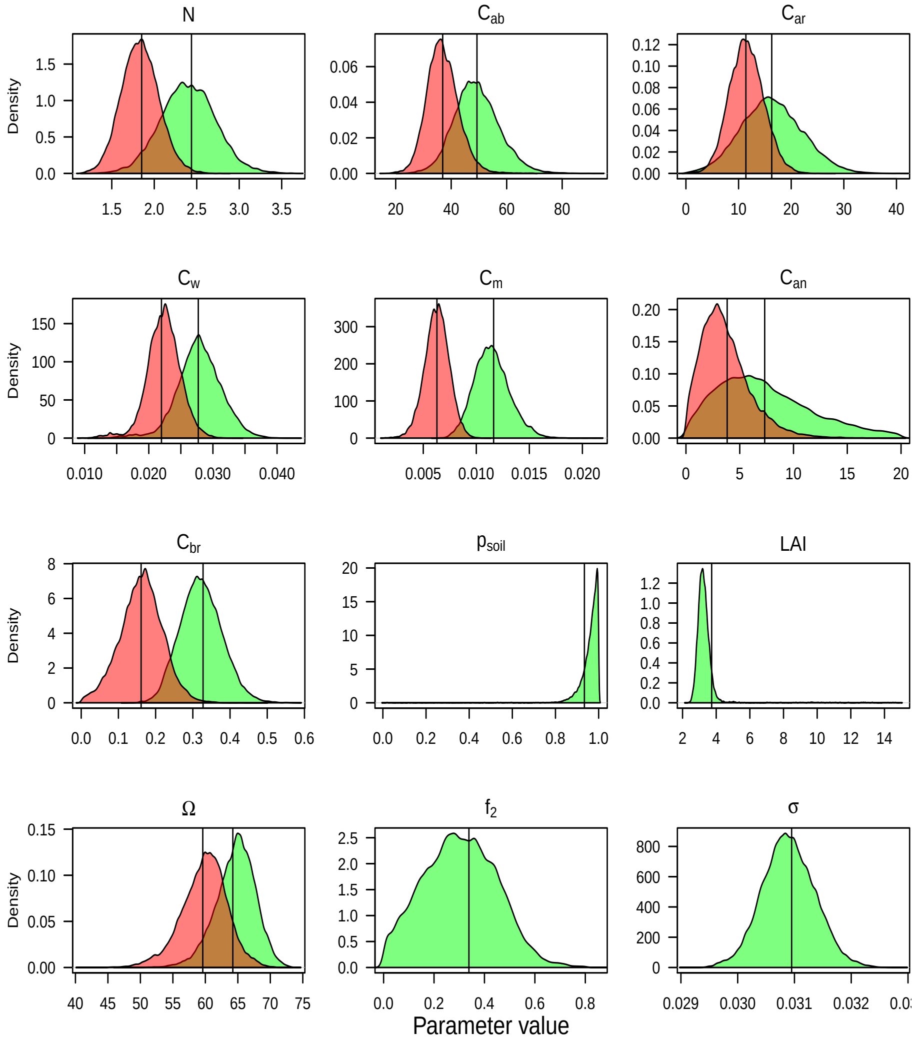


Figure S5.3: Marginal density plots for the posterior distributions for each model parameter of the PROSAIL2 model fit to the Panama dataset. Parameters that differ between lianas and trees are given in red and green respectively. Vertical lines indicate mean values for all distributions.

### S6 Absolute Validation

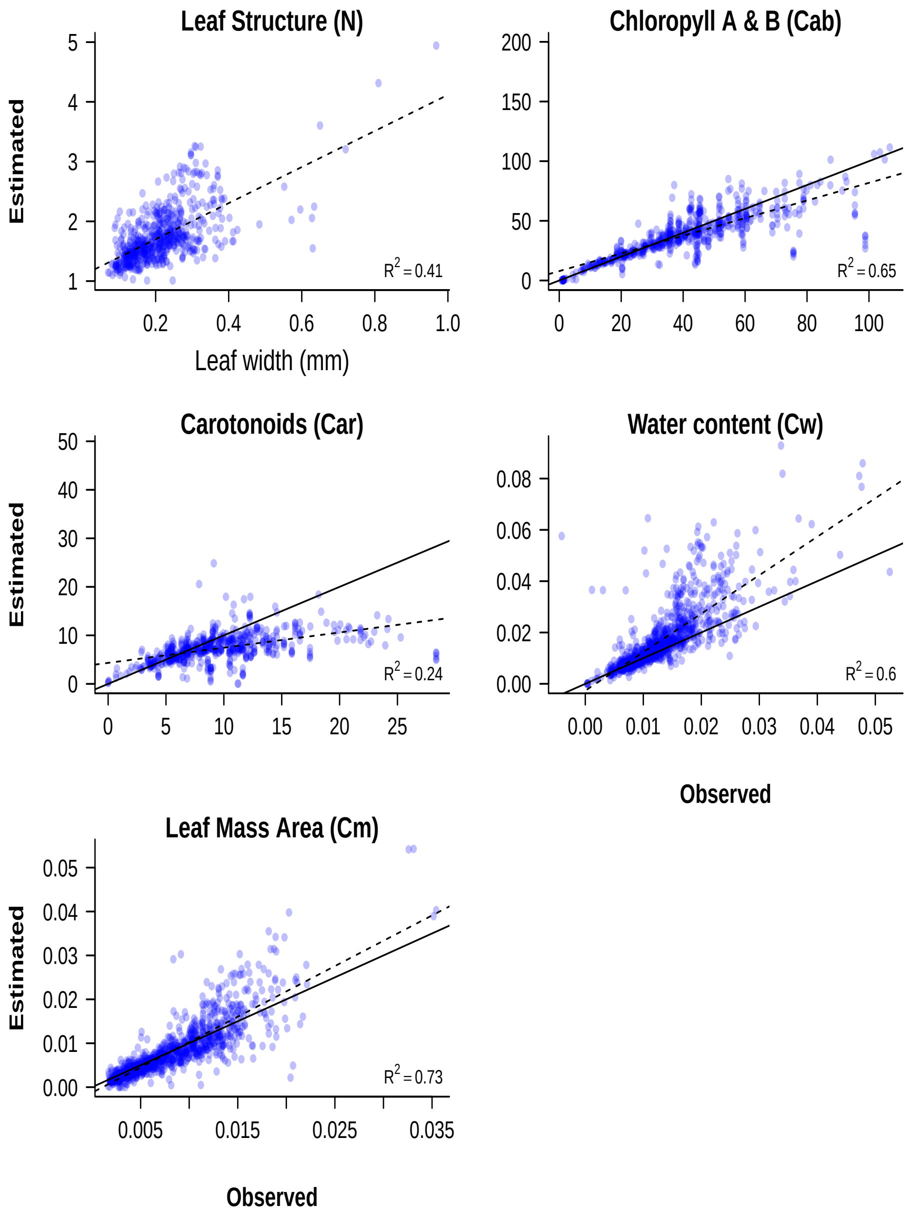


Figure S6.1: Leaf scale model inversion results compared to independent ground estimates for PROSPECT5B. Dashed lines show the linear relationship between x and y, as estimated by OLS. Thick black line is the 1:1 line. Note that all plots represent the same traits, and have the same units, with exception of the first panel.

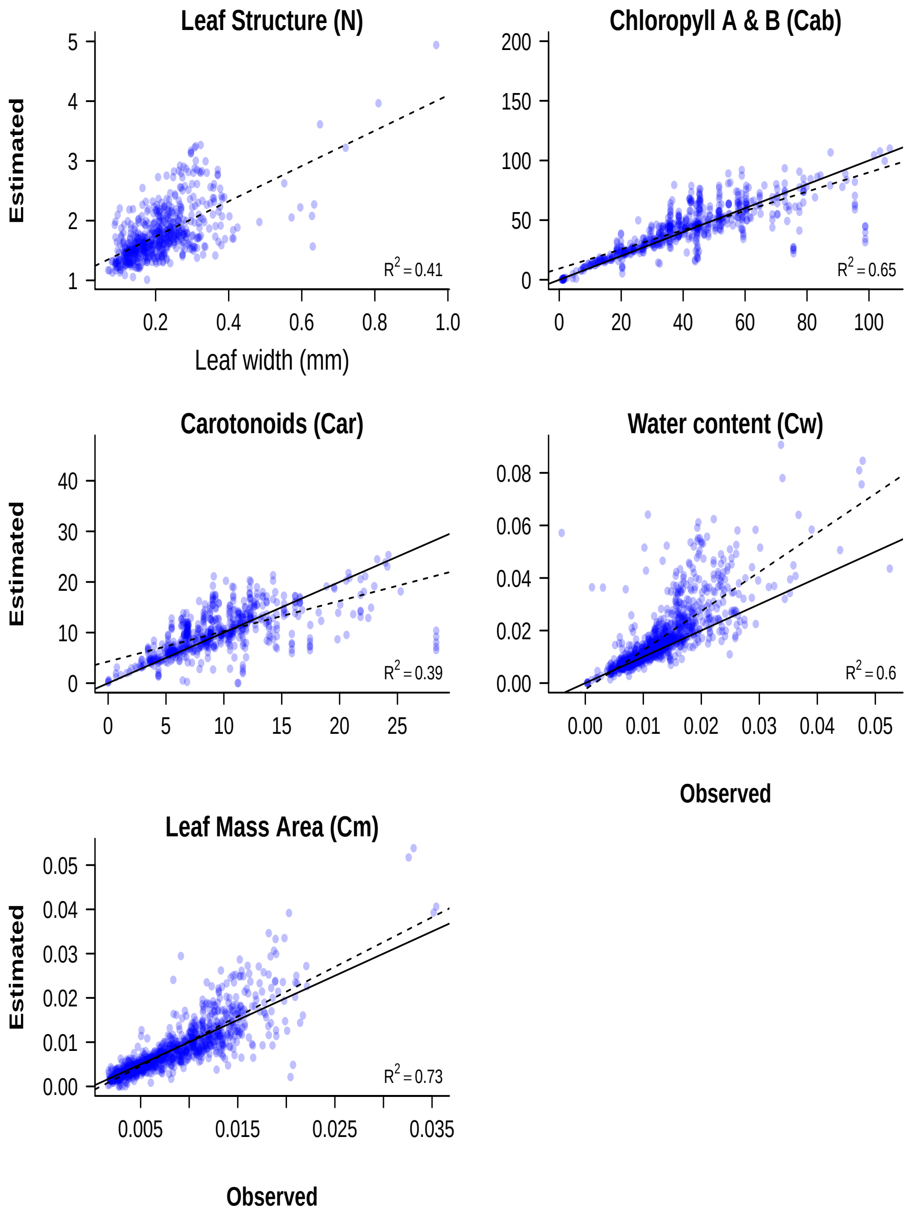


Figure S6.2: Leaf scale model inversion results compared to independent ground estimates for PROSPECTD. Dashed lines show the linear relationship between x and y, as estimated by OLS. Thick black line is the 1:1 line. Note that all plots represent the same traits, and have the same units, with exception of the first panel.

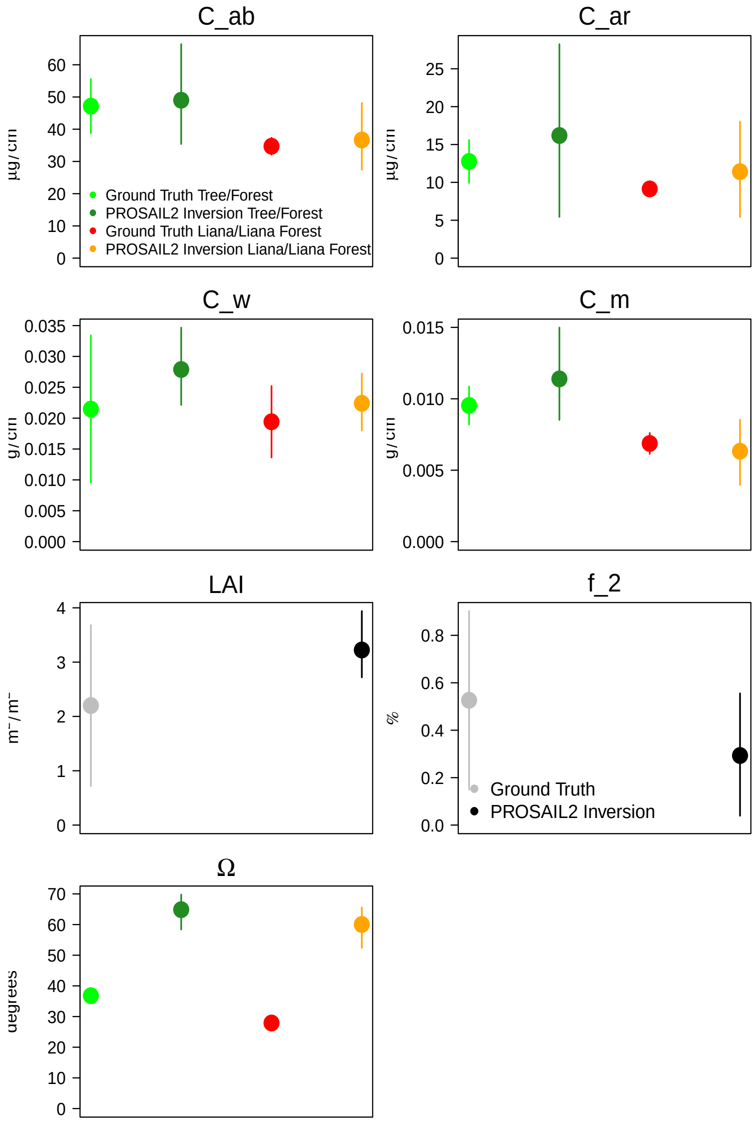


Figure S6.3: Canopy scale model inversion results for Panama (for Gigante) compared to independent ground estimates (from San Lorenzo, Barro Colorado Island, and PNM). Parameters that differed between lianas and trees are given in shades of red and green respectively. LAI and $f_2$ are given in shades of grey (see legends).

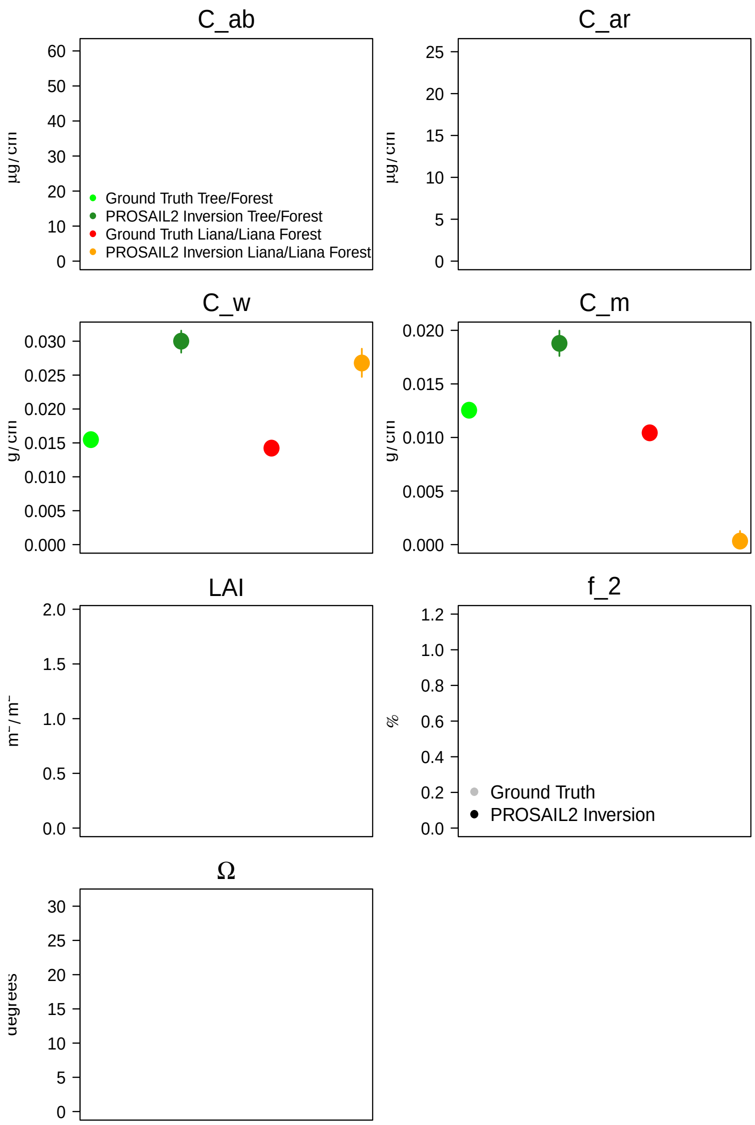


Figure S6.4: Canopy scale model inversion results for Malaysia (Danum Valley) compared to independent ground estimates (Sepilok). Parameters that differed between lianas and trees are given in shades of red and green respectively. LAI and $f_2$ are given in shades of grey (see legends). Blank plots represent traits with no ground 1validation for this study site.

### S7 Leaf profiles

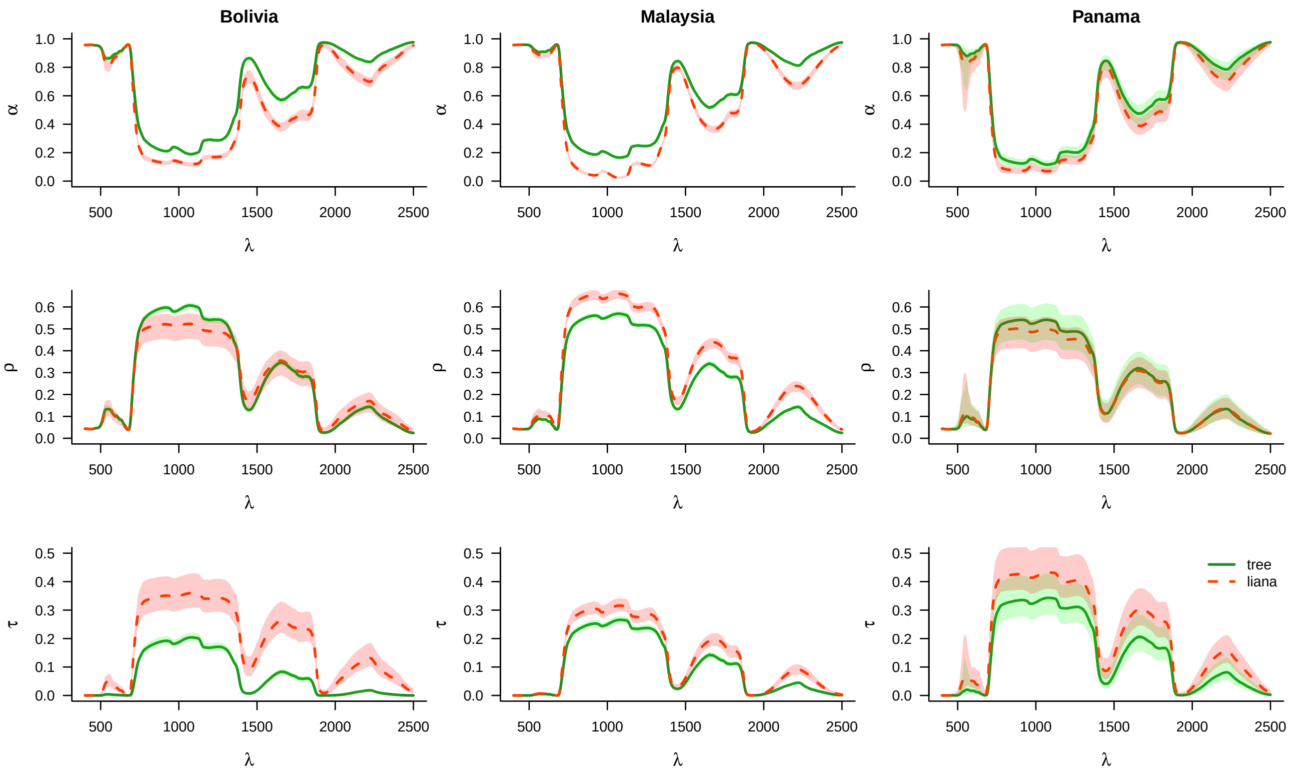


Figure S7.1: Leaf absorbtion ($\alpha$), reflectance ($\rho$) and transmission ($\tau$) profiles against wavelength ($\lambda$) for trees (green) and lianas (red) - based on the inverse PROSAIL2 fits for reflectance data collected in Bolivia, Malaysia, and Panama.

#### S8 Model experiment per site

This is the elaboration of figure 6 from the maintext towards individual sites.

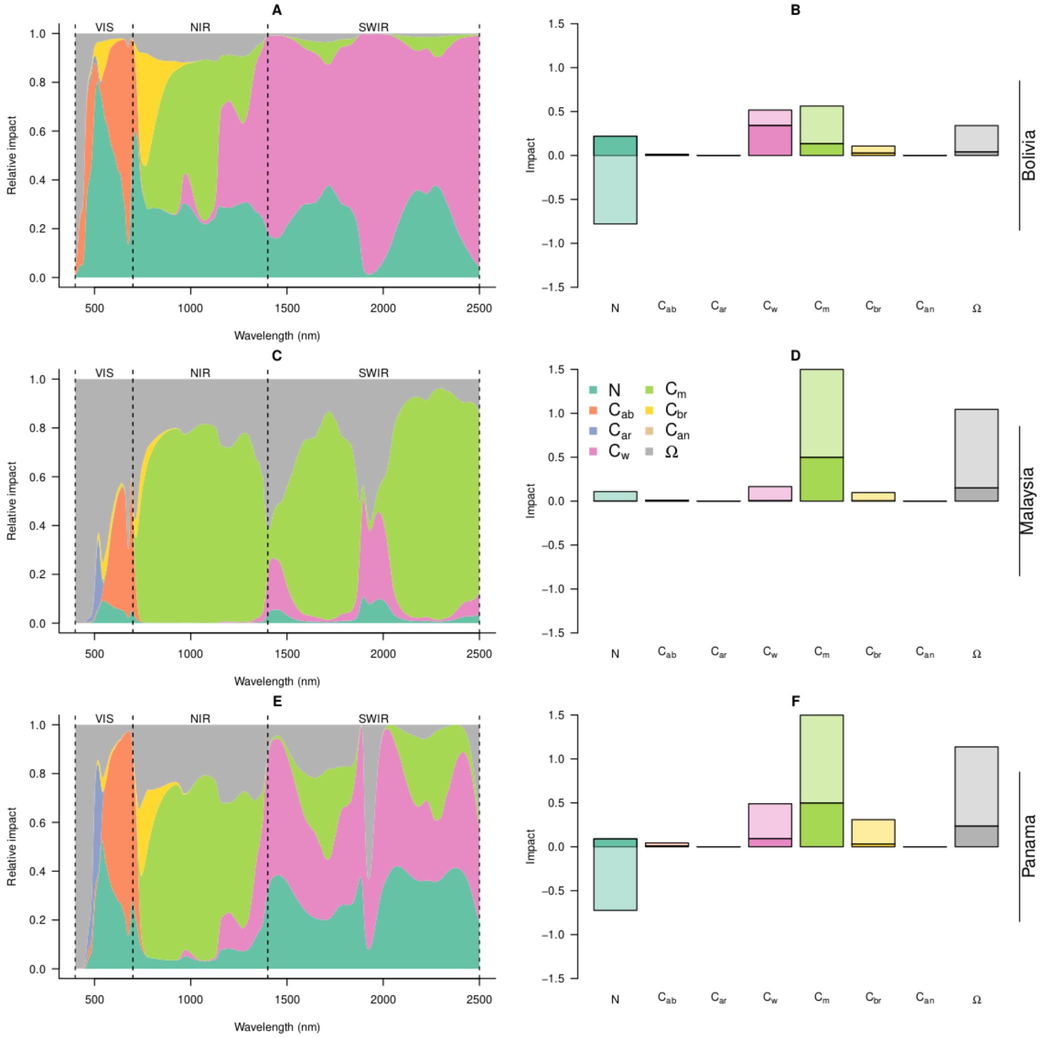


Figure S8.1: Model experiment showing the relative importance of different traits in generating the liana signal. In all model experiments, the traits in the top layer were set to tree traits and then iteratively replaced with liana traits either one by one (additive) or in pairs (interactive effects). At each interaction, the change in the Kullback-Leiber divergence score was calculated. Panels A, C, and E give the relative absolute impact of changing a single trait in the top canopy layer at each wavelength. Panels B, D and F give the total impact of additive (darker colors) and interactive effects (lighter colors). Here, a positive impact refers to a change in the direction of the liana signal, while a negative impact refers to in a change away from the liana signal.

### S9 Multiple scattering

For specular flux, the energy that scatters after an interaction with a leaf particle can be roughly approximated for lianas as $(\tau_1 + \rho_1)^N$ and trees as $(\tau_2 + \rho_2)^N$ - where N gives the number of successive interactions with a leaf particles. The rate at which the transmitted energy decays (i.e. is absorbed) is then given by the derivative of the above functions: $log((\tau_1 + \rho_1))(\tau_1 + \rho_1)^N$ and $log((\tau_2 + \rho_2))(\tau_2 + \rho_2)^N$ respectively. It follows that when $(\tau_1 + \rho_1) > (\tau_2 + \rho_2)$, the energy declines at a slower rate for lianas with each interation N (as seen below in panel A), which generally will lead to more energy being available for a secondary scattering including reflection towards the observer. The average rate of energy decline is generally much lower for lianas across the entire solar spectrum (see panel B).

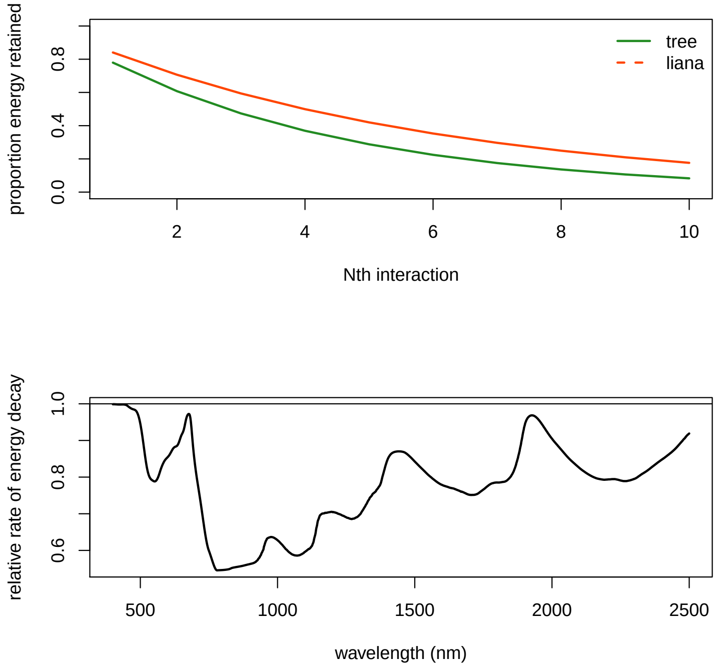


Figure S9.1: Top panel: The proportion of energy expected available for multiple scattering events at a wavelength of 1299 nm for trees (green) and lianas (red) based on the inverse PROSAIL2 fits for Panama (A). Bottom panel: The relative rate of decline for liana leaves vs tree leaves at the first interaction (N=1) - which can be up to 55% lower for lianas, a value of 1 means that energy declines equally for interactions with liana vs tree leaves, a value $<1$ incates it declines slower for lianas.
